# Identification of potential Auxin Response Candidate genes for soybean rapid canopy coverage through comparative evolution and expression analysis

**DOI:** 10.1101/2023.10.26.564213

**Authors:** Deisiany F. Neres, Joseph S. Taylor, John A. Bryant, Bastiaan O. R. Bargmann, R. Clay Wright

## Abstract

*Glycine max*, soybean, is an abundantly cultivated crop worldwide. Efforts have been made over the past decades to improve soybean production in traditional and organic agriculture, driven by growing demand for soybean-based products. Rapid canopy cover development (RCC) increases soybean yields and suppresses early-season weeds.

Genome-wide association studies have found natural variants associated with RCC, however causal mechanisms are unclear. Auxin modulates plant growth and development and has been implicated in RCC traits. Therefore, modulation of auxin regulatory genes may enhance RCC. Here, we focus on the use of genomic tools and existing datasets to identify auxin signaling pathway RCC candidate genes, using a comparative phylogenetics and expression analysis approach.

We identified genes encoding 14 TIR1/AFB auxin receptors, 61 Aux/IAA auxin co-receptors and transcriptional co-repressors, and 55 ARF auxin response factors in the soybean genome. We used Bayesian phylogenetic inference to identify soybean orthologs of *Arabidopsis thaliana* genes, and defined an ortholog naming system for these genes. To further define potential auxin signaling candidate genes for RCC, we examined tissue-level expression of these genes in existing datasets and identified highly expressed auxin signaling genes in apical tissues early in development. We identified at least 4 *TIR1/AFB*, 8 *Aux/IAA*, and 8 *ARF* genes with highly specific expression in one or more RCC-associated tissues. We hypothesize that modulating the function of these genes through gene editing or traditional breeding will have the highest likelihood of affecting RCC while minimizing pleiotropic effects.

## Introduction

Soybean [*Glycine max* (L.) Merr.] is one of the most cultivated crops worldwide. Soybeans are a chief source of plant-based protein and are commonly used in animal feed, dairy, fuel, and oil production. Due to increased demand for soybean-based products for both traditional and organic agriculture, great effort has been made over the decades to improve soybean production. However, when it comes to organic agriculture, weeds are still a significant impediment to producers (Horn *et al*. 1985; Wang *et al*. 2014; Leinonen et al., 2019; Lusk, 2022). Crop-weed interference studies have shown that soybean yield loss can reach up to 90% if necessary management practices aren’t in place. Furthermore, the impact on soybean grain yield is dependent on weed species and density (Horn *et al*., 1985; Silva *et al*., 2009).

Rapid canopy cover development (RCC) is a highly valuable trait for soybean, as it enables early-season weed suppression and allows the crop plant to outcompete and shade weeds (Peters *et al.,* 1965; Horn *et al*. 1985; Xavier *et al., 2017*). Several studies have found that the trait of RCC has a positive effect on soybean yields, especially in organic production conditions. For example, Xavier *et al*. (2017) investigated the genetic architecture of RCC and found that this trait is associated with higher grain yields in soybeans (r = 0.87). Additionally, Peters *et al*. (1965) observed that RCC reduced weed biomass and increased soybean yields in a row spacing experiment. These findings suggest that RCC is an important trait for improving soybean yields, and may be particularly advantageous in organic production where weed competition is a challenge.

RCC is primarily related to plant aerial architecture, which encompasses a range of structures including hypocotyl, cotyledon, apical and axillary meristematic tissues, and leaves. By providing greater available leaf area sooner after planting, plants with improved RCC can increase solar radiation interception, which is crucial for photosynthesis and ultimately dictates crop growth and yield (Stewart *et al*. 2003; Edwards *et al.,* 2005; Hatfield *et al.,* 2019). Additionally, increased radiation interception by the desired crop plant will shade weeds, potentially inhibiting their germination and growth. Soybean plants that exhibit RCC also benefit from improved water-use efficiency by minimizing water evaporation and enhancing soil moisture retention (Peters *et al.,* 1965). RCC can help improve soybean production and weed management, making this an important area of research for agricultural systems. Despite the potential advantages of RCC, the causal mechanisms for this developmental trait are not yet fully understood.

Soybean Genome-Wide Association Studies (GWAS) have shown that auxin is important in early establishment of canopy cover (Xavier *et al., 2017;* Kaler *et al.,* 2018; Li *et al*., 2023). For instance, among seven SNPs significantly associated with RCC, two are found in a locus that contains three auxin related genes (Xavier et al., 2017). Additionally, Kaler *et al*. (2018) identified 92 RCC-correlated SNPs, two of which are directly auxin related, and several more which are auxin responsive. Li *et al*. (2023) hypothesized that *AtARF7* ortholog is involved in RCC. Therefore, auxin related genes are a potential RCC breeding target and worth exploring further. Auxin is a phytohormone involved in numerous aspects of plant growth and development, including response to biotic and abiotic stresses (Padmanabhan *et al*., 2005; Sun *et al.,* 2016), root and seed development, apical dominance (Tatematsu *et al*., 2004; Prigge *et al*., 2020), leaf longevity and expansion, and plant architecture (Davies, 1995; Lim *et al*., 2010). Auxin signaling genes in soybean have previously been associated with root nodulation and development, as well as flowering (Sun *et al.,* 2016; Cai *et al.,* 2017; Li and Chen, 2023). Due to the complexity of the auxin signaling network and its interaction with other signaling pathways, little is known about how these genes affect the aerial architecture of soybean plants (Davies, 1995; Swarup *et al*., 2002; Vernoux *et al*., 2011; Calderón-Villalobos *et al*., 2012; Lavy and Estelle, 2016).

The auxin transcriptional regulatory system is comprised of three main gene families: the receptors *TRANSPORT INHIBITOR RESPONSE 1/ AUXIN SIGNALING F-BOX (TIR1/AFB)*, the transcriptional repressors *AUXIN/INDOLE-3-ACETIC ACID INDUCIBLE (Aux/IAA)*, and transcription factors *AUXIN RESPONSE FACTOR (ARF)* genes. When auxin levels in a plant cell are low, Aux/IAA transcriptional repressors are bound to ARF transcription factors repressing auxin-responsive gene expression through Aux/IAA interaction with TOPLESS/TOPLESS-RELATED (TPL/TPR) co-repressor proteins (Abel *et al*., 1996; Tiwari *et al.,* 2001; Overvoorde *et al.,* 2005; Weijers *et al*. 2005; Szemenyei *et al.,* 2008). When auxin accumulates, it acts as a molecular glue increasing the affinity between the members of the TIR1/AFB *receptors* and Aux/IAA repressors, which form auxin co-receptor complexes (Tan *et al*.. 2007). Ultimately, as the majority of TIR1/AFB proteins associate with SKP1-CULLIN-F-box ubiquitin ligase complex, the Aux/IAAs bound to these complex are subjected to polyubiquitination targeting them for proteolysis through the 26S proteasome. Degradation of the Aux/IAAs leads to de-repression of activator ARFs and expression of auxin responsive genes (Gray *et al*., 2001; Ramos *et al.,* 2001; Zenser *et al*., 2001; Chapman and Estelle, 2009). Interplay between these three auxin gene families varies in a tissue-dependent manner and is responsible for orchestrating different plant fate and agronomic traits.

Several auxin signaling genes are known to play many important roles in *Arabidopsis* apical meristem development that point to auxin’s involvement in RCC. Perhaps the most notable are the *ARF1/2* and *ARF3/4* clades of repressor ARFs. Mutants in *ARF2* have enlarged rosette leaves and seeds as well as elongated hypocotyls, but at the cost of reduced fertility (Okushima *et al*., 2005a). *arf1/arf2* double mutants have even stronger developmental phenotypes (Okushima *et al*., 2005b). Variants affecting *ARF3/ETTIN* yield pleiotropic effects on leaf and flower development as well as abnormal phyllotaxy (Nishimura *et al*., 2005; Pecker *et al*., 2005). *ARF3* and *ARF4* are regulated by trans-acting siRNAs which, when disrupted, lead to changes in the progressive leaf shape from round, flat juvenile leaves to oblong, downward curling (epinastic) adult leaves, also known as heteroblasty. Consistent with this association between relief of auxin response gene repression and increased growth of aerial tissues, mutants in the activators *ARF6* and *ARF8* result in dwarfing of aerial tissues (Nagpal *et al*., 2005; Okushima *et al*., 2005b). These ARFs are similarly small-RNA-regulated, in this case by miRNA167. Additionally, *arf7/arf19* double mutant plants are of small stature (Okushima *et al*., 2005b).

In Arabidopsis, auxin perception via TIR1/AFB–auxin–Aux/IAA interaction is less clearly associated with RCC related traits than some *arf* phenotypes noted above. *TIR1/AFB* and *Aux/IAA* mutants are more commonly associated with root traits such as formation of lateral roots, but still there is evidence for their regulation of apical dominance and meristematic tissues, as well as leaf longevity and other above-ground developmental processes (Dharmasiri *et al*., 2005; Parry *et al*., 2009; Lim *et al*., 2010; Salehin *et al*., 2015). The TIR1/AFB auxin receptor F-box genes have overlapping functions and are expressed and accumulated in growing organs related to RCC in Arabidopsis, such as shoot apical meristem (SAM) and leaf primordia (Parry *et al*. 2009). Mutants in two or more of the TIR1/AFB genes of Arabidopsis become increasingly dwarfed (Prigge *et al*., 2020). In the Arabidopsis Aux/IAA family there is more evidence for tissue-specificity in expression and function (Overvoorde *et al*., 2005). For instance, IAA3 preferentially regulates ARF7 and ARF19 during root development, whereas during leaf expansion and hypocotyl tropic responses these same response factors are modulated by IAA19 and IAA28 (Wilmoth *et al*., 2005). However, loss-of-function mutants in *Aux/IAA* genes show subtle or no phenotypes, likely due to redundancy or compensation within this large gene family (Nagpal *et al*., 2000; Tian *et al*., 2002; Overvoorde *et al*., 2005).

The existing knowledge of auxin perception and related traits in Arabidopsis and other model species can be useful in identifying gene functions in crop species such as maize and soybean. A similar approach to ours was used to examine the evolutionary developmental relationships between *Arabidopsis* and *Zea maize* auxin signaling components (Matthes *et al*., 2019). Moreover, auxin genes have been linked to shoot height in soybean plants, such as up-regulation of *GmIAA9* and *GmIAA29* leading to internode elongation, *GmARF9* promoting first pod height, and a dwarf phenotype being associated with *GmIAA27* (Jiang *et al*., 2018; Su *et al*., 2022; Zhang *et al*., 2022). Additionally, other genes associated with both gibberellin and cytokinin signaling pathways, have also been reported to play a role in shoot architecture development in soybean, such as CYTOKININ RESPONSE FACTOR *4a* (*GmCRF4a*). In previous phylogenetic studies, YUCCA flavin monooxygenases, encoded by *YUC* genes, have been pointed out to be in control of auxin biosynthesis in Arabidopsis, *Oryza sativa*, *Medicago truncatula, Lotus japonicus,* and *Glycine max* (Wang *et al*., 2019)(reviewed in Li and Chen, 2023). Furthermore, auxin biosynthesis is tightly regulated by feedback through the auxin perception pathway (Cao *et al*., 2019).

Here, we report findings from phylogenetics coupled with transcriptomic analysis to predict genes associated with RCC development in soybean. We identified orthologous groups of *TIR1/AFB*, *Aux/IAA*, and *ARF* genes from *Arabidopsis* and several *Fabaceae* species, and defined an ortholog-based naming system for these *Glycine max* genes to help facilitate evolutionary comparisons. We then performed an expression analysis based on our hypothesis that these auxin signaling genes that are highly and specifically expressed in apical tissues and early development will have the greatest effect on RCC development (RCC-related tissues). Auxin signaling genes are responsible for many developmental processes in plants, thus engineering RCC through manipulation of auxin signaling can be perilous due to pleiotropic effects on plant growth and development. To avoid this pitfall, we propose to target genes associated with specific RCC tissues as identified via principal component analysis. As a result of phylogenetic and transcriptomic analysis, we identified several candidate auxin-signaling genes that potentially affect RCC. Soybean orthologs of *Arabidopsis ARF2, ARF8,* and *ARF9* were found to be expressed specifically in RCC-related tissues. We also identified a selection of *Aux/IAA* and *TIR1/AFB* candidate genes. Notably, *Aux/IAA* genes revealed a higher tissue specificity in soybean expression datasets, corroborating with the existing body of literature of spatial expression analysis in other species. These findings suggest promising RCC candidate genes for further exploration at both the molecular and organismal levels.

## Results

### Expression analysis and identification of auxin signaling targets for RCC development in *G. max*

First, we present our expression analysis using PCA with gene names presented as ortholog names from our phylogenetic analysis, below. Expression data retrieved from both NCBI SRA repository and SoyBase (defined and referenced in the “Expression analysis data” section in methodology) indicate that a total of 133 of 221 transcripts for these combined gene families displayed median expression equal or greater than 2 TPM across RCC-related tissues. The 133 transcripts were further evaluated through PCA in order to identify which auxin regulatory genes are most specifically associated with certain RCC tissues (Figure 1). The first two principal components (PCs) account for 66.5% and 15.2%, respectively, of the total variation in gene expression. Therefore, the two-dimensional scatter-plot of the 133 given transcripts shown in figure 1 represents 81.7% of total variation.

**Figure 1.**
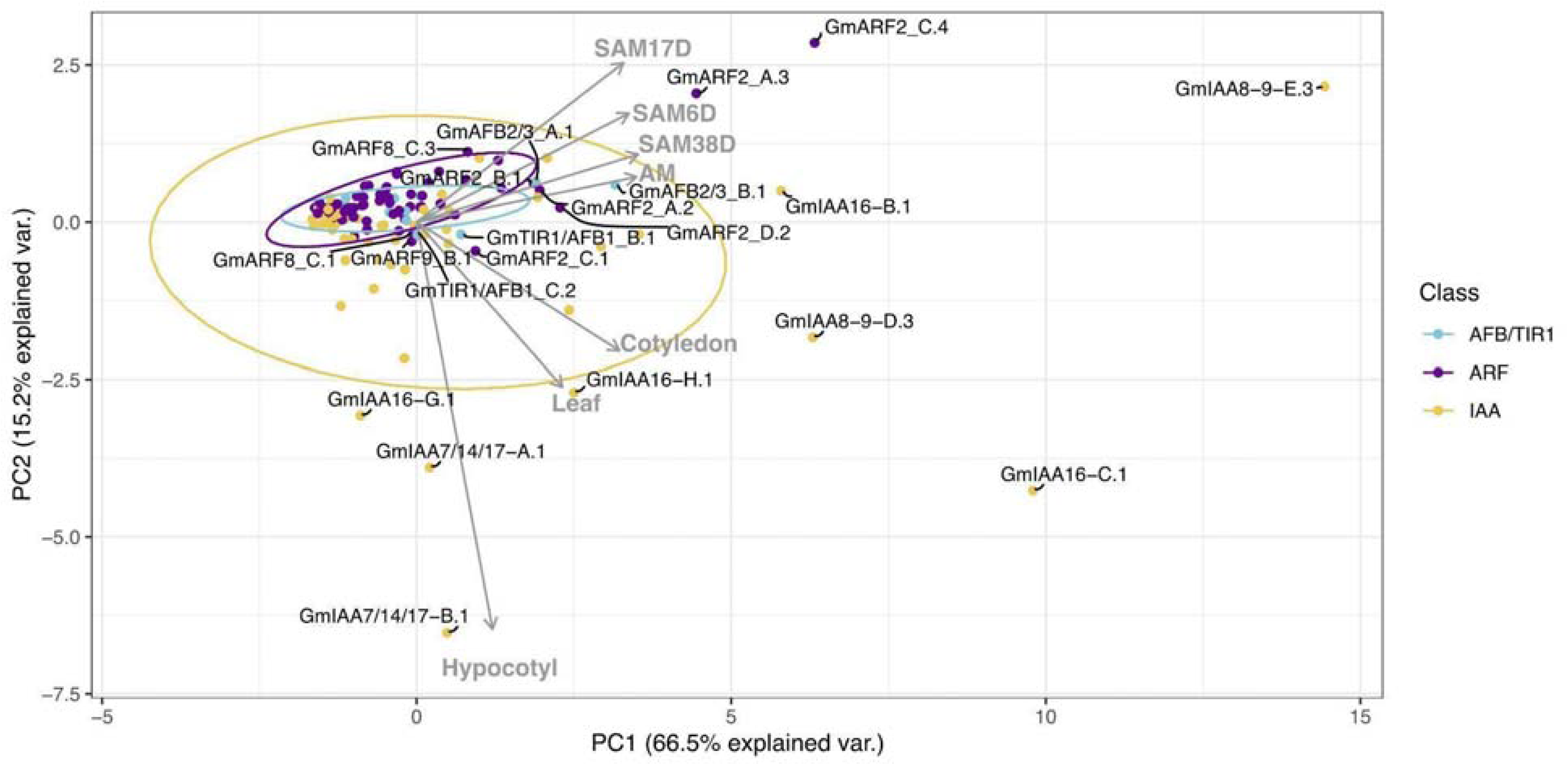
Correlation-based Principal Component Analysis (PCA): biplot of gene transcript expression and explanatory tissues involved in plant aerial architecture as eigenvectors (grey arrows), n = 7). Principal components 1 and 2 account for 81.7% of the total inertia. Ellipses are used here as a visual representation of dispersion of data points within each group (TIR1/AFB, ARF, and Aux/IAA (IAA)) with a 70% confidence interval. TIR1/AFB genes are colored cyan, ARF genes are colored purple, and Aux/IAA (IAA) genes are colored yellow. Some labels are connected to their respective points with hard lines. Genes clustering together inside the ellipses are hypothesized to have more pleiotropic effects on plant growth and development, whereas genes associated with a specific RCC tissue (genes that fall along an eigenvector, outside of the respectively colored ellipse) are hypothesized to have narrower effects and be more amenable to engineering RCC traits through gene editing.

Correlation between gene expression patterns in different tissues can be qualitatively assessed based on the angle and distance between their eigenvectors (grey arrows in Figure 1). Expression levels across meristematic tissues were strongly correlated and primary contributors to PC1 (Figure 1). Gene expression levels in hypocotyl, leaf, and cotyledon tissues were also positively correlated with one another, although less strongly so, and contribute more to PC2. The relationship between hypocotyl and meristematic tissues ranges from no correlation to a weak negative correlation, probably due to *Aux/IAAs* tissue specificity. We observe a 90 degree angle between hypocotyl and axillary meristem (AM) tissues, representing no correlation between gene expression in these tissues. To assess the variance and outliers in expression for each of these gene families, ellipses representing 70% confidence levels, assuming a Student’s T-distribution, in each PC for each gene family were drawn. Genes within these ellipses (in yellow, purple, and blue), indicate that most genes do not show strong tissue-specific expression, as has been previously shown for these gene families in other plant species (Matthes *et al*., 2019). Therefore, genes clustered within ellipses, and close to the origin are more likely to display pleiotropic effects. This is suggested by the ellipses’ overlap in our PCA space, as overlapping ellipses imply a high similarity between the expression of these genes and the tissues under analysis. Additionally, genes positioned closer to the origin are thought to make minor contributions to the variance explained by a principal component. It is worth noting that this could also be a result of the low overall expression levels, which further reduces their impact on the analysis.

Variability in gene expression specificity across different gene families can be inferred from the area and shape of their respective ellipses. Our observations reveal that Aux/IAAs (yellow) exhibit a larger ellipse with approximately equal width along both PC1 and PC2 axes. This configuration signifies higher variability in expression between tissues within the Aux/IAA gene family. In contrast, ARFs (purple) and TIR1/AFBs (blue) display smaller, more elongated ellipses, particularly towards meristematic tissues. This suggests that these genes are less variable and possibly share more functional overlap. Collectively, these findings align with the existing body of literature, underscoring the idea that the regulation of auxin response may be tissue-dependent, primarily influenced by Aux/IAAs repressor proteins.

Similarly, tissue specificity can be qualitatively assessed based on the placement of genes with respect to these eigenvectors. We defined genes falling outside the ellipses as having strong association with one or more tissues, as indicated by the eigenvectors of tissue expression also shown on the PC biplots. Auxin regulatory genes strongly associated with meristematic tissues were *GmAFB2/3_A.1/B.1*, *GmARF2_A.2/A.3/C.4/D.2*, *GmARF8_C.3, GmIAA8-9_E.3*, and *GmIAA16_B.1* (Figure 1). There were five Aux/IAAs (*GmIAA7/14/17_A.1/B.1, IAA16_G.1/H.1*), five ARFs (*GmARF9_B.1, ARF2_C.5,* and *ARF8_C.1*), and two AFBs (*GmTIR1/AFB1_B.1* and *C.2*) associated with hypocotyl, leaf, and cotyledon tissues (Figure - 1). *IAA8-9_D.3, IAA16_C.1,* and *ARF2_C1* are highly expressed in both leaf, cotyledon and meristematic tissues, as they fall between these eigenvectors. We also observed that PC2 and PC3 (10.7% explained variance) provide additional information and other possible candidate gene contributions to these aerial tissues (Figure S1). With *GmARF9_B.1*, and *ARF2_C.1* contributing to leaf and cotyledon development, as well as *GmARF11/18_A.2* to meristematic tissues (Figure S1).

We have also compared our principal component analysis results using all 14 tissues from the databases to those using only the 7 tissues that are part of aerial architecture. It is evident that our principal component analysis is minimally affected by the exclusion of root, callus and later development tissues (Figure S2). Therefore, we are confident in displaying only the aerial tissues that are of interest to us. Additionally, we have observed that the root, hypocotyl, nodule, and open flower tissues align along the same direction, suggesting a correlation between these tissues. As we delve into principal components that explain smaller variations, we notice an enhanced discrimination in the correlation between root, nodule, and hypocotyl tissues (Figure S3). Further investigation of these genes is essential to gain more insightful information about the genes that play a crucial role in the growth and development of these tissues. Additionally, it will help us determine if different pairs of auxin regulatory genes are significant contributors to RCC.

### Phylogenetic analysis of TIR1/AFB co-receptors and proposed orthology based on *A. thaliana* classification

We propose a nomenclature for *G. max* auxin signaling genes using a comparative phylogenetic approach with *A. thaliana*. This nomenclature aims to enhance the prediction of gene and protein functions in *G. max* by leveraging the extensive gene function knowledge available in *A. thaliana*. Several evolutionary and developmental comparative studies have shown that genes that share sequence similarity, and therefore fall within the same clade on a phylogeny, are broadly predicted to have similar function (Zhou *et al*., 2013; Hyung *et al*., 2014; Husbands *et al*., 2015; Damodharan *et al*., 2018; Zhang *et al*., 2021). Although comparative approaches have been used to identify and predict *G. max* gene function, little is known about the role of auxin signaling in *G. max* aerial architecture. In *A. thaliana* several auxin signaling genes are associated with unique aerial phenotypes of their mutants (Husbands *et al*., 2015; Zhou *et al*., 2013; Damodharan *et al*., 2018; Zhang *et al.,* 2021; Parry *et al*., 2009; Dharmasiri *et al*., 2005). Therefore, we have assigned ortholog names for each *G. max* auxin signaling gene based on their phylogenetic placement in clades with *A. thaliana* genes, e.g. *GmTIR1/AFB1_A* is more closely related to (shares more sequence similarity with) *AtTIR1* and *AtAFB1* than other members of the *A. thaliana TIR1/AFB* family. Furthermore, *G. max* has undergone two whole-genome duplications. The first corresponds to the early legume-duplication, which occurred approximately 59 million years (Myr) ago. The second duplication is Glycine-specific and happened around 13 Myr ago (Schmutz *et al*., 2010). As a result, *G. max* typically possesses more than one copy of each *A. thaliana* ortholog (Figures 2, 4, and 5). Because of this, we have assigned letters to differentiate between *G. max* orthologs for each clade. Additionally, to strengthen our classifications of these *G. max* gene families, and provide some additional context of these clades in the *Fabaceae*, we have also included other well studied legume species *G. soja*, *M. truncatula*, and *L. japonicus*.

**Figure 2.**
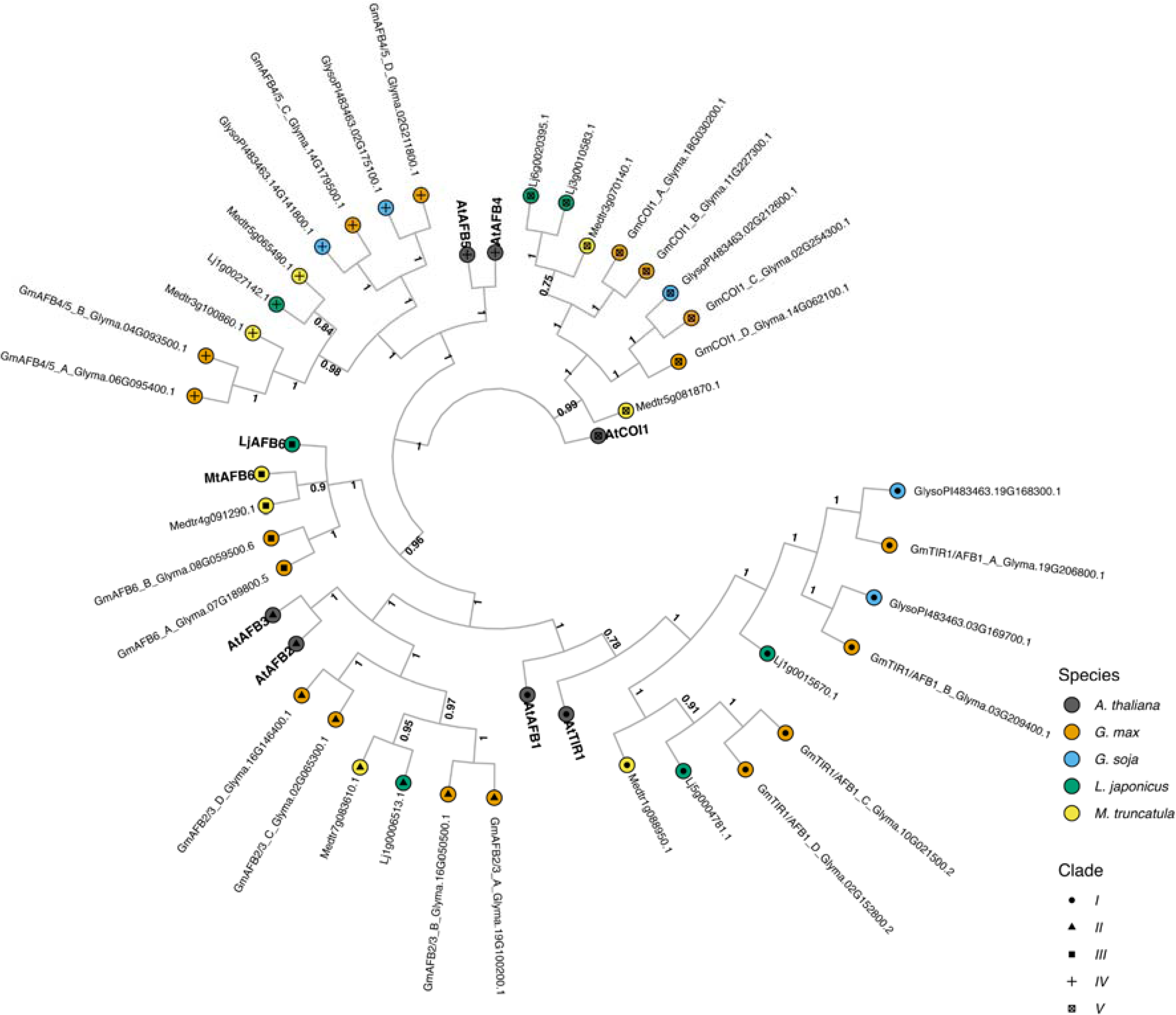
The evolutionary relationships between *G. max* TIR1/AFB proteins, and *A. thaliana* and other legume species orthologs. The historical relationship was inferred using the Bayesian Inference method (Ronquist *et al*., 2012). The optimal tree is drawn according to the posterior probability of the evolutionary distances. The posterior probability of each node is labeled. Each tip is colored according to species, with *A. thaliana* in black, *G. max* in orange, *G. soja* in light blue, *L. japonicus* in green, and *M. truncatula* in yellow. *A. thaliana* gene symbols are displayed in bold to better visualize clade separation. Clades are also defined by the shapes towards tip extremities, with clade I as a circle, clade II as a triangle, clade III as a square, clade IV as an addition sign, and clade V as a ballot box with an x. *G. max* genes were named according to their orthology to *A. thaliana* followed by their gene ID.

The fourteen *G. max* TIR1/AFB auxin receptors identified were grouped into five main clades. Clade I, which has the least similarity to the COI1 jasmonate receptor outgroup, comprises *A. thaliana* TIR1-like proteins, with a posterior probability of 0.78 (the lowest probability found in the resulting tree) to its *G. soja*, *M. truncatula*, *L. japonicus,* and *G. max* orthologs. This clade consists of four *G. max* proteins GmTIR1/AFB1_A–D, of which only two (GmTIR1/AFB1_A and B) have a *G. soja* sister taxa, the other sister taxa were lost in *G. soja* (Figure 2). Interestingly this A/B subclade contains a *L. japonicus* taxa but is missing a *M. truncatula* taxa, suggesting a complex pattern of recent gene loss events in this clade. Clade II, consists of AFB2/AFB3-like proteins and also contains four *G. max* representatives but does not contain any *G. soja* and only one representative each from *M. truncatula* and *L. japonicus* (Figure 2). Clade III, is comprised by the AFB6-like proteins, Medtr8g098695.2 and Lj4g0012889.1 defined in Rogato *et al*. (2021), and, according to Parry *et al*. (2009), this clade was lost during the evolution of the Brassicaceae (*A. thaliana* family) and Poaceae families. We identified two *G. max* AFB6 orthologs. Interestingly, *G. soja*, *G. max*’s wild relative, lacks orthologs for both clades II and III, whereas *M. truncatula* and *L. japonicus* are both represented. Clade IV, AFB4/AFB5-like proteins, contains four *G. max* proteins, of which again only two have a corresponding *G. soja* sister as with TIR1/AFB1-like clade I. Lastly, clade V, comprises COI1-like (Coronatine Insensitive) F-box proteins, with four *G. max* homologous proteins and only one representative sister in *G. soja*. The difference in number of genes between cultivated soybean, *G. max*, and its wild relative, *G. soja* is also in accordance with comparative genomics published data in which cultivated soybean has many unique genes that are unavailable in its wild relative (Joshi *et al*., 2013 and references therein). Moreover, all the *G. max* orthologs here identified contain all the necessary functional domains to perceive auxin and associate with the SCF E3 ligase complex necessary for the turnover of transcriptional repressors and auxin mediated response (Figure S4 and Appendix A).

### Phylogenetic analysis of Aux/IAA co-receptors and proposed orthology based on *A. thaliana* classification

Relative to the TIR1/AFB family, both the Aux/IAA and ARF families are far more numerous in all plants. Additionally, both have been more broadly studied in previous phylogenetic analysis. However the names assigned to *G. max* genes/proteins are often not defined by orthology, making comparative inference difficult. To facilitate such analysis here and in the future we have assigned *G. max* gene names for these families according to their orthology to *A. thaliana* (Table 1).

**Table 1.** Full list of *G. max* orthologous genes with Gene Identities, its assigned orthology, transcript identifier, classification, assigned clade and lastly its assigned family.

| Gene ID | Orthology | Transcript ID | Class | Clade | Family |
| --- | --- | --- | --- | --- | --- |
| Glyma.19G206800 | GmTIR1/AFB1_A.1 | Glyma.19G206800.1 | TIR1/AFB1 | I | AFB/TIR1 |
| Glyma.03G209400 | GmTIR1/AFB1_B.1 | Glyma.03G209400.1 | TIR1/AFB1 | I | AFB/TIR1 |
| Glyma.10G021500 | GmTIR1/AFB1_C.1 | Glyma.10G021500.1 | TIR1/AFB1 | I | AFB/TIR1 |
| Glyma.10G021500 | GmTIR1/AFB1_C.2 | Glyma.10G021500.2 | TIR1/AFB1 | I | AFB/TIR1 |
| Glyma.10G021500 | GmTIR1/AFB1_C.3 | Glyma.10G021500.3 | TIR1/AFB1 | I | AFB/TIR1 |
| Glyma.02G152800 | GmTIR1/AFB1_D.1 | Glyma.02G152800.1 | TIR1/AFB1 | I | AFB/TIR1 |
| Glyma.02G152800 | GmTIR1/AFB1_D.2 | Glyma.02G152800.2 | TIR1/AFB1 | I | AFB/TIR1 |
| Glyma.19G100200 | GmAFB2/3_A.1 | Glyma.19G100200.1 | AFB2/3 | II | AFB/TIR1 |
| Glyma.16G050500 | GmAFB2/3_B.1 | Glyma.16G050500.1 | AFB2/3 | II | AFB/TIR1 |
| Glyma.02G065300 | GmAFB2/3_C.1 | Glyma.02G065300.1 | AFB2/3 | II | AFB/TIR1 |
| Glyma.16G146400 | GmAFB2/3_D.1 | Glyma.16G146400.1 | AFB2/3 | II | AFB/TIR1 |
| Glyma.07G189800 | GmAFB6_A.3 | Glyma.07G189800.3 | AFB6 | III | AFB/TIR1 |
| Glyma.07G189800 | GmAFB6_A.4 | Glyma.07G189800.4 | AFB6 | III | AFB/TIR1 |
| Glyma.07G189800 | GmAFB6_A.5 | Glyma.07G189800.5 | AFB6 | III | AFB/TIR1 |
| Glyma.08G059500 | GmAFB6_B.6 | Glyma.08G059500.6 | AFB6 | III | AFB/TIR1 |
| Glyma.06G095400 | GmAFB4/5_A.1 | Glyma.06G095400.1 | AFB4/5 | IV | AFB/TIR1 |
| Glyma.04G093500 | GmAFB4/5_B.1 | Glyma.04G093500.1 | AFB4/5 | IV | AFB/TIR1 |
| Glyma.14G179500 | GmAFB4/5_C.1 | Glyma.14G179500.1 | AFB4/5 | IV | AFB/TIR1 |
| Glyma.02G211800 | GmAFB4/5_D.1 | Glyma.02G211800.1 | AFB4/5 | IV | AFB/TIR1 |
| Glyma.18G030200 | GmCOI1_A.1 | Glyma.18G030200.1 | COI1 | V | AFB/TIR1 |
| Glyma.11G227300 | GmCOI1_B.1 | Glyma.11G227300.1 | COI1 | V | AFB/TIR1 |
| Glyma.16G023600 | GmARF11/18_A.1 | Glyma.16G023600.1 | B | I | ARF |
| Glyma.16G023600 | GmARF11/18_A.2 | Glyma.16G023600.2 | B | I | ARF |
| Glyma.16G023600 | GmARF11/18_A.3 | Glyma.16G023600.3 | B | I | ARF |
| Glyma.07G054800 | GmARF11/18_B.1 | Glyma.07G054800.1 | B | I | ARF |
| Glyma.07G054800 | GmARF11/18_B.2 | Glyma.07G054800.2 | B | I | ARF |
| Glyma.07G054800 | GmARF11/18_B.3 | Glyma.07G054800.3 | B | I | ARF |
| Glyma.07G054800 | GmARF11/18_B.4 | Glyma.07G054800.4 | B | I | ARF |
| Glyma.03G258300 | GmARF11/18_C.1 | Glyma.03G258300.1 | B | I | ARF |
| Glyma.15G142600 | GmARF13_A.1 | Glyma.15G142600.1 | B | I | ARF |
| Glyma.15G142600 | GmARF13_A.2 | Glyma.15G142600.2 | B | I | ARF |
| Glyma.15G142600 | GmARF13_A.3 | Glyma.15G142600.3 | B | I | ARF |
| Glyma.12G164100 | GmARF1_A.1 | Glyma.12G164100.1 | B | I | ARF |
| Glyma.16G000300 | GmARF1_B.1 | Glyma.16G000300.1 | B | I | ARF |
| Glyma.07G272800 | GmARF1_C.1 | Glyma.07G272800.1 | B | I | ARF |
| Glyma.06G164900 | GmARF2_A.2 | Glyma.06G164900.2 | B | I | ARF |
| Glyma.06G164900 | GmARF2_A.3 | Glyma.06G164900.3 | B | I | ARF |
| Glyma.04G200600 | GmARF2_B.1 | Glyma.04G200600.1 | B | I | ARF |
| Glyma.04G200600 | GmARF2_B.2 | Glyma.04G200600.2 | B | I | ARF |
| Glyma.05G200800 | GmARF2_C.1 | Glyma.05G200800.1 | B | I | ARF |
| Glyma.05G200800 | GmARF2_C.4 | Glyma.05G200800.4 | B | I | ARF |
| Glyma.05G200800 | GmARF2_C.5 | Glyma.05G200800.5 | B | I | ARF |
| Glyma.08G008100 | GmARF2_D.2 | Glyma.08G008100.2 | B | I | ARF |
| Glyma.08G008100 | GmARF2_D.3 | Glyma.08G008100.3 | B | I | ARF |
| Glyma.03G208800 | GmARF2_E.1 | Glyma.03G208800.1 | B | I | ARF |
| Glyma.07G202200 | GmARF3_A.1 | Glyma.07G202200.1 | B | I | ARF |
| Glyma.13G174000 | GmARF3_B.1 | Glyma.13G174000.1 | B | I | ARF |
| Glyma.13G174000 | GmARF3_B.2 | Glyma.13G174000.2 | B | I | ARF |
| Glyma.13G174000 | GmARF3_B.3 | Glyma.13G174000.3 | B | I | ARF |
| Glyma.13G234200 | GmARF3_C.1 | Glyma.13G234200.1 | B | I | ARF |
| Glyma.12G071000 | GmARF4_A.1 | Glyma.12G071000.1 | B | I | ARF |
| Glyma.12G071000 | GmARF4_A.2 | Glyma.12G071000.2 | B | I | ARF |
| Glyma.12G171000 | GmARF4_C.1 | Glyma.12G171000.1 | B | I | ARF |
| Glyma.12G171000 | GmARF4_C.2 | Glyma.12G171000.2 | B | I | ARF |
| Glyma.12G171000 | GmARF4_C.3 | Glyma.12G171000.3 | B | I | ARF |
| Glyma.12G171000 | GmARF4_C.4 | Glyma.12G171000.4 | B | I | ARF |
| Glyma.12G171000 | GmARF4_C.5 | Glyma.12G171000.5 | B | I | ARF |
| Glyma.12G171000 | GmARF4_C.6 | Glyma.12G171000.6 | B | I | ARF |
| Glyma.12G171000 | GmARF4_C.7 | Glyma.12G171000.7 | B | I | ARF |
| Glyma.12G171000 | GmARF4_C.8 | Glyma.12G171000.8 | B | I | ARF |
| Glyma.12G171000 | GmARF4_C.9 | Glyma.12G171000.9 | B | I | ARF |
| Glyma.01G103500 | GmARF9_A.1 | Glyma.01G103500.1 | B | I | ARF |
| Glyma.01G103500 | GmARF9_A.2 | Glyma.01G103500.2 | B | I | ARF |
| Glyma.03G070500 | GmARF9_B.1 | Glyma.03G070500.1 | B | I | ARF |
| Glyma.03G070500 | GmARF9_B.2 | Glyma.03G070500.2 | B | I | ARF |
| Glyma.07G134800 | GmARF9_C.1 | Glyma.07G134800.1 | B | I | ARF |
| Glyma.18G184500 | GmARF9_D.1 | Glyma.18G184500.1 | B | I | ARF |
| Glyma.18G184500 | GmARF9_D.2 | Glyma.18G184500.2 | B | I | ARF |
| Glyma.18G184500 | GmARF9_D.3 | Glyma.18G184500.3 | B | I | ARF |
| Glyma.14G217700 | GmARF5_A.1 | Glyma.14G217700.1 | A | II | ARF |
| Glyma.17G256500 | GmARF5_B.1 | Glyma.17G256500.1 | A | II | ARF |
| Glyma.08G100100 | GmARF6_A.2 | Glyma.08G100100.2 | A | II | ARF |
| Glyma.08G100100 | GmARF6_A.3 | Glyma.08G100100.3 | A | II | ARF |
| Glyma.13G221400 | GmARF6_C.1 | Glyma.13G221400.1 | A | II | ARF |
| Glyma.13G221400 | GmARF6_C.2 | Glyma.13G221400.2 | A | II | ARF |
| Glyma.15G091000 | GmARF6_D.1 | Glyma.15G091000.1 | A | II | ARF |
| Glyma.15G091000 | GmARF6_D.2 | Glyma.15G091000.2 | A | II | ARF |
| Glyma.15G091000 | GmARF6_D.3 | Glyma.15G091000.3 | A | II | ARF |
| Glyma.15G091000 | GmARF6_D.4 | Glyma.15G091000.4 | A | II | ARF |
| Glyma.02G281700 | GmARF6_E.1 | Glyma.02G281700.1 | A | II | ARF |
| Glyma.02G281700 | GmARF6_E.2 | Glyma.02G281700.2 | A | II | ARF |
| Glyma.02G281700 | GmARF6_E.3 | Glyma.02G281700.3 | A | II | ARF |
| Glyma.14G032700 | GmARF6_F.1 | Glyma.14G032700.1 | A | II | ARF |
| Glyma.14G032700 | GmARF6_F.2 | Glyma.14G032700.2 | A | II | ARF |
| Glyma.09G072200 | GmARF7/19_A.1 | Glyma.09G072200.1 | A | II | ARF |
| Glyma.15G181000 | GmARF7/19_B.1 | Glyma.15G181000.1 | A | II | ARF |
| Glyma.15G181000 | GmARF7/19_B.2 | Glyma.15G181000.2 | A | II | ARF |
| Glyma.17G047100 | GmARF7/19_C.1 | Glyma.17G047100.1 | A | II | ARF |
| Glyma.13G112600 | GmARF7/19_D.1 | Glyma.13G112600.1 | A | II | ARF |
| Glyma.07G130400 | GmARF7/19_E.1 | Glyma.07G130400.1 | A | II | ARF |
| Glyma.07G130400 | GmARF7/19_E.2 | Glyma.07G130400.2 | A | II | ARF |
| Glyma.07G130400 | GmARF7/19_E.3 | Glyma.07G130400.3 | A | II | ARF |
| Glyma.01G002100 | GmARF7/19_F.1 | Glyma.01G002100.1 | A | II | ARF |
| Glyma.01G002100 | GmARF7/19_F.2 | Glyma.01G002100.2 | A | II | ARF |
| Glyma.05G221300 | GmARF7/19_G.1 | Glyma.05G221300.1 | A | II | ARF |
| Glyma.05G221300 | GmARF7/19_G.2 | Glyma.05G221300.2 | A | II | ARF |
| Glyma.05G221300 | GmARF7/19_G.3 | Glyma.05G221300.3 | A | II | ARF |
| Glyma.08G027800 | GmARF7/19_H.1 | Glyma.08G027800.1 | A | II | ARF |
| Glyma.08G027800 | GmARF7/19_H.2 | Glyma.08G027800.2 | A | II | ARF |
| Glyma.11G204200 | GmARF8_A.1 | Glyma.11G204200.1 | A | II | ARF |
| Glyma.11G204200 | GmARF8_A.2 | Glyma.11G204200.2 | A | II | ARF |
| Glyma.18G046800 | GmARF8_B.1 | Glyma.18G046800.1 | A | II | ARF |
| Glyma.02G239600 | GmARF8_C.1 | Glyma.02G239600.1 | A | II | ARF |
| Glyma.02G239600 | GmARF8_C.2 | Glyma.02G239600.2 | A | II | ARF |
| Glyma.02G239600 | GmARF8_C.3 | Glyma.02G239600.3 | A | II | ARF |
| Glyma.02G239600 | GmARF8_C.4 | Glyma.02G239600.4 | A | II | ARF |
| Glyma.02G239600 | GmARF8_C.5 | Glyma.02G239600.5 | A | II | ARF |
| Glyma.14G208500 | GmARF8_D.1 | Glyma.14G208500.1 | A | II | ARF |
| Glyma.12G174100 | GmARF10/16_A.1 | Glyma.12G174100.1 | C | III | ARF |
| Glyma.13G325200 | GmARF10/16_B.1 | Glyma.13G325200.1 | C | III | ARF |
| Glyma.12G076200 | GmARF10/16_C.1 | Glyma.12G076200.1 | C | III | ARF |
| Glyma.11G145500 | GmARF10/16_D.1 | Glyma.11G145500.1 | C | III | ARF |
| Glyma.10G210600 | GmARF10/16_E.1 | Glyma.10G210600.1 | C | III | ARF |
| Glyma.20G180000 | GmARF10/16_F.1 | Glyma.20G180000.1 | C | III | ARF |
| Glyma.10G053500 | GmARF17_A.1 | Glyma.10G053500.1 | C | III | ARF |
| Glyma.13G140600 | GmARF17_B.1 | Glyma.13G140600.1 | C | III | ARF |
| Glyma.13G140600 | GmARF17_B.2 | Glyma.13G140600.2 | C | III | ARF |
| Glyma.19G181900 | GmARF17_C.2 | Glyma.19G181900.2 | C | III | ARF |
| Glyma.13G084700 | GmARF17_D.1 | Glyma.13G084700.1 | C | III | ARF |
| Glyma.04G254200 | GmARF17_F.1 | Glyma.04G254200.1 | C | III | ARF |
| Glyma.19G168500 | GmIAA12-13-A.1 | Glyma.19G168500.1 | A | I | IAA |
| Glyma.19G168500 | GmIAA12-13-A.2 | Glyma.19G168500.2 | A | I | IAA |
| Glyma.19G168500 | GmIAA12-13-A.3 | Glyma.19G168500.3 | A | I | IAA |
| Glyma.19G168500 | GmIAA12-13-A.4 | Glyma.19G168500.4 | A | I | IAA |
| Glyma.19G168500 | GmIAA12-13-A.5 | Glyma.19G168500.5 | A | I | IAA |
| Glyma.19G168500 | GmIAA12-13-A.6 | Glyma.19G168500.6 | A | I | IAA |
| Glyma.03G167400 | GmIAA12-13-B.10 | Glyma.03G167400.10 | A | I | IAA |
| Glyma.03G167400 | GmIAA12-13-B.11 | Glyma.03G167400.11 | A | I | IAA |
| Glyma.03G167400 | GmIAA12-13-B.9 | Glyma.03G167400.9 | A | I | IAA |
| Glyma.13G127000 | GmIAA12-13-C.3 | Glyma.13G127000.3 | A | I | IAA |
| Glyma.10G040400 | GmIAA12-13-D.3 | Glyma.10G040400.3 | A | I | IAA |
| Glyma.07G015200 | GmIAA18/26/28-A.1 | Glyma.07G015200.1 | A | I | IAA |
| Glyma.08G200700 | GmIAA18/26/28-B.1 | Glyma.08G200700.1 | A | I | IAA |
| Glyma.08G200700 | GmIAA18/26/28-B.2 | Glyma.08G200700.2 | A | I | IAA |
| Glyma.13G354100 | GmIAA18/26/28-C.11 | Glyma.13G354100.11 | A | I | IAA |
| Glyma.13G354100 | GmIAA18/26/28-C.6 | Glyma.13G354100.6 | A | I | IAA |
| Glyma.13G354100 | GmIAA18/26/28-C.8 | Glyma.13G354100.8 | A | I | IAA |
| Glyma.15G020300 | GmIAA18/26/28-D.1 | Glyma.15G020300.1 | A | I | IAA |
| Glyma.15G020300 | GmIAA18/26/28-D.2 | Glyma.15G020300.2 | A | I | IAA |
| Glyma.05G229300 | GmIAA27-A.1 | Glyma.05G229300.1 | A | I | IAA |
| Glyma.01G039300 | GmIAA27-B.2 | Glyma.01G039300.2 | A | I | IAA |
| Glyma.08G036400 | GmIAA27-C.1 | Glyma.08G036400.1 | A | I | IAA |
| Glyma.08G036400 | GmIAA27-C.2 | Glyma.08G036400.2 | A | I | IAA |
| Glyma.09G193000 | GmIAA27-D.1 | Glyma.09G193000.1 | A | I | IAA |
| Glyma.13G356600 | GmIAA27-E.1 | Glyma.13G356600.1 | A | I | IAA |
| Glyma.15G017500 | GmIAA27-F.1 | Glyma.15G017500.1 | A | I | IAA |
| Glyma.15G017500 | GmIAA27-F.2 | Glyma.15G017500.2 | A | I | IAA |
| Glyma.15G017500 | GmIAA27-F.3 | Glyma.15G017500.3 | A | I | IAA |
| Glyma.07G018100 | GmIAA27-G.1 | Glyma.07G018100.1 | A | I | IAA |
| Glyma.08G203100 | GmIAA27-H.1 | Glyma.08G203100.1 | A | I | IAA |
| Glyma.08G203100 | GmIAA27-H.2 | Glyma.08G203100.2 | A | I | IAA |
| Glyma.06G067700 | GmIAA29-A.1 | Glyma.06G067700.1 | A | I | IAA |
| Glyma.04G066300 | GmIAA29-B.1 | Glyma.04G066300.1 | A | I | IAA |
| Glyma.13G159000 | GmIAA29-C.1 | Glyma.13G159000.1 | A | I | IAA |
| Glyma.17G112300 | GmIAA29-D.1 | Glyma.17G112300.1 | A | I | IAA |
| Glyma.10G270500 | GmIAA32/34-A.1 | Glyma.10G270500.1 | A | I | IAA |
| Glyma.20G120800 | GmIAA32/34-B.1 | Glyma.20G120800.1 | A | I | IAA |
| Glyma.09G203300 | GmIAA8-9-A.1 | Glyma.09G203300.1 | A | I | IAA |
| Glyma.09G203300 | GmIAA8-9-A.2 | Glyma.09G203300.2 | A | I | IAA |
| Glyma.09G203300 | GmIAA8-9-A.3 | Glyma.09G203300.3 | A | I | IAA |
| Glyma.01G019400 | GmIAA8-9-B.1 | Glyma.01G019400.1 | A | I | IAA |
| Glyma.01G019400 | GmIAA8-9-B.2 | Glyma.01G019400.2 | A | I | IAA |
| Glyma.01G019400 | GmIAA8-9-B.3 | Glyma.01G019400.3 | A | I | IAA |
| Glyma.01G019400 | GmIAA8-9-B.4 | Glyma.01G019400.4 | A | I | IAA |
| Glyma.08G273500 | GmIAA8-9-C.1 | Glyma.08G273500.1 | A | I | IAA |
| Glyma.08G273500 | GmIAA8-9-C.2 | Glyma.08G273500.2 | A | I | IAA |
| Glyma.01G098000 | GmIAA8-9-D.3 | Glyma.01G098000.3 | A | I | IAA |
| Glyma.06G091700 | GmIAA8-9-E.3 | Glyma.06G091700.3 | A | I | IAA |
| Glyma.04G089900 | GmIAA8-9-F.1 | Glyma.04G089900.1 | A | I | IAA |
| Glyma.14G185400 | GmIAA8-9-G.1 | Glyma.14G185400.1 | A | I | IAA |
| Glyma.14G185400 | GmIAA8-9-G.2 | Glyma.14G185400.2 | A | I | IAA |
| Glyma.02G218100 | GmIAA8-9-H.2 | Glyma.02G218100.2 | A | I | IAA |
| Glyma.02G218100 | GmIAA8-9-H.4 | Glyma.02G218100.4 | A | I | IAA |
| Glyma.10G180000 | GmIAA1-4-A.1 | Glyma.10G180000.1 | B | II | IAA |
| Glyma.10G180000 | GmIAA1-4-A.2 | Glyma.10G180000.2 | B | II | IAA |
| Glyma.20G210500 | GmIAA1-4-B.1 | Glyma.20G210500.1 | B | II | IAA |
| Glyma.10G000700 | GmIAA1-4-C.2 | Glyma.10G000700.2 | B | II | IAA |
| Glyma.02G000500 | GmIAA1-4-D.1 | Glyma.02G000500.1 | B | II | IAA |
| Glyma.19G161000 | GmIAA1-4-E.1 | Glyma.19G161000.1 | B | II | IAA |
| Glyma.19G161000 | GmIAA1-4-E.2 | Glyma.19G161000.2 | B | II | IAA |
| Glyma.19G161000 | GmIAA1-4-E.3 | Glyma.19G161000.3 | B | II | IAA |
| Glyma.03G158600 | GmIAA1-4-F.1 | Glyma.03G158600.1 | B | II | IAA |
| Glyma.10G031800 | GmIAA1-4-G.3 | Glyma.10G031800.3 | B | II | IAA |
| Glyma.02G142600 | GmIAA1-4-H.1 | Glyma.02G142600.1 | B | II | IAA |
| Glyma.02G142600 | GmIAA1-4-H.2 | Glyma.02G142600.2 | B | II | IAA |
| Glyma.17G042800 | GmIAA31-A.1 | Glyma.17G042800.1 | B | II | IAA |
| Glyma.17G042800 | GmIAA31-A.2 | Glyma.17G042800.2 | B | II | IAA |
| Glyma.13G117100 | GmIAA31-B.1 | Glyma.13G117100.1 | B | II | IAA |
| Glyma.13G117100 | GmIAA31-B.2 | Glyma.13G117100.2 | B | II | IAA |
| Glyma.13G117100 | GmIAA31-B.3 | Glyma.13G117100.3 | B | II | IAA |
| Glyma.13G117100 | GmIAA31-B.4 | Glyma.13G117100.4 | B | II | IAA |
| Glyma.19G221900 | GmIAA31-C.1 | Glyma.19G221900.1 | B | II | IAA |
| Glyma.19G221900 | GmIAA31-C.2 | Glyma.19G221900.2 | B | II | IAA |
| Glyma.03G224800 | GmIAA31-D.1 | Glyma.03G224800.1 | B | II | IAA |
| Glyma.10G138500 | GmIAA31-E.1 | Glyma.10G138500.1 | B | II | IAA |
| Glyma.02G007300 | GmIAA31-F.1 | Glyma.02G007300.1 | B | II | IAA |
| Glyma.13G361100 | GmIAA6/19-A.1 | Glyma.13G361100.1 | B | II | IAA |
| Glyma.15G012800 | GmIAA6/19-B.1 | Glyma.15G012800.1 | B | II | IAA |
| Glyma.07G034200 | GmIAA6/19-C.1 | Glyma.07G034200.1 | B | II | IAA |
| Glyma.13G361200 | GmIAA15-A.2 | Glyma.13G361200.2 | C | III | IAA |
| Glyma.13G361200 | GmIAA15-A.3 | Glyma.13G361200.3 | C | III | IAA |
| Glyma.13G361200 | GmIAA15-A.4 | Glyma.13G361200.4 | C | III | IAA |
| Glyma.13G361200 | GmIAA15-A.5 | Glyma.13G361200.5 | C | III | IAA |
| Glyma.15G012700 | GmIAA15-B.1 | Glyma.15G012700.1 | C | III | IAA |
| Glyma.15G012700 | GmIAA15-B.2 | Glyma.15G012700.2 | C | III | IAA |
| Glyma.15G012700 | GmIAA15-B.3 | Glyma.15G012700.3 | C | III | IAA |
| Glyma.15G012700 | GmIAA15-B.4 | Glyma.15G012700.4 | C | III | IAA |
| Glyma.15G012700 | GmIAA15-B.5 | Glyma.15G012700.5 | C | III | IAA |
| Glyma.10G162400 | GmIAA16-A.1 | Glyma.10G162400.1 | C | III | IAA |
| Glyma.10G162400 | GmIAA16-A.2 | Glyma.10G162400.2 | C | III | IAA |
| Glyma.10G162400 | GmIAA16-A.3 | Glyma.10G162400.3 | C | III | IAA |
| Glyma.20G225000 | GmIAA16-B.1 | Glyma.20G225000.1 | C | III | IAA |
| Glyma.03G247400 | GmIAA16-C.1 | Glyma.03G247400.1 | C | III | IAA |
| Glyma.19G245200 | GmIAA16-D.1 | Glyma.19G245200.1 | C | III | IAA |
| Glyma.10G031900 | GmIAA16-E.1 | Glyma.10G031900.1 | C | III | IAA |
| Glyma.10G031900 | GmIAA16-E.2 | Glyma.10G031900.2 | C | III | IAA |
| Glyma.02G142500 | GmIAA16-F.3 | Glyma.02G142500.3 | C | III | IAA |
| Glyma.19G161100 | GmIAA16-G.1 | Glyma.19G161100.1 | C | III | IAA |
| Glyma.03G158700 | GmIAA16-H.1 | Glyma.03G158700.1 | C | III | IAA |
| Glyma.10G180100 | GmIAA7/14/17-A.1 | Glyma.10G180100.1 | C | III | IAA |
| Glyma.20G210400 | GmIAA7/14/17-B.1 | Glyma.20G210400.1 | C | III | IAA |

The *G. max Aux/IAA* gene family contains sixty-one members. They are classified according to our bootstrapping support into three main classes A, B and C which are also divided into Clade I, II, and III, respectively (Figure 3). Most of the sixty-one Aux/IAAs fall in class A followed by B and C. All PB1, EAR motif, and degron domains are conserved in protein coding sequence except for orthologs of AtIAA15/20/29/30/32/33/34 which lack one or more of these domains (Appendix B). Clade I, is composed by orthologs of AtIAA8/9/10/11/12/13/18/26/27/28/32/34, with at least two representatives of *G. max*. In contrast, its wild relative, *G. soja*, has one representative sister for each GmIAA10-13, but lacks at least one sister to the remaining *G. max* orthologous groups. Clade II, constituted of AtIAA1/2/3/4/5/6/19/20/30/31 orthologs, is distributed similarly to Clade I, however all orthologous groups are missing at least one sister in *G. soja*. Clade III, is composed by AtIAA7/14/15/16/17/33 orthologs as well as the root of our phylogeny. There are two *G. max* orthologs of AtIAA7/14/17, and each of them has an ortholog in *G. soja*. In contrast, orthologs of AtIAA15 and 33, have only one *G. max* ortholog. The GmIAA15 has one *G. soja* ortholog, but GmIAA33 does not have a *G. soja* ortholog. Although most species do have a representative, due to its paleopolyploidy, *G. max* has the most abundant number of Aux/IAAs when compared to other species analyzed here. In particular, the IAA16 orthology group contains eight *G. max* orthologs in two subclades of four, one sharing a more recent common ancestor with AtIAA16.

**Figure 3.**
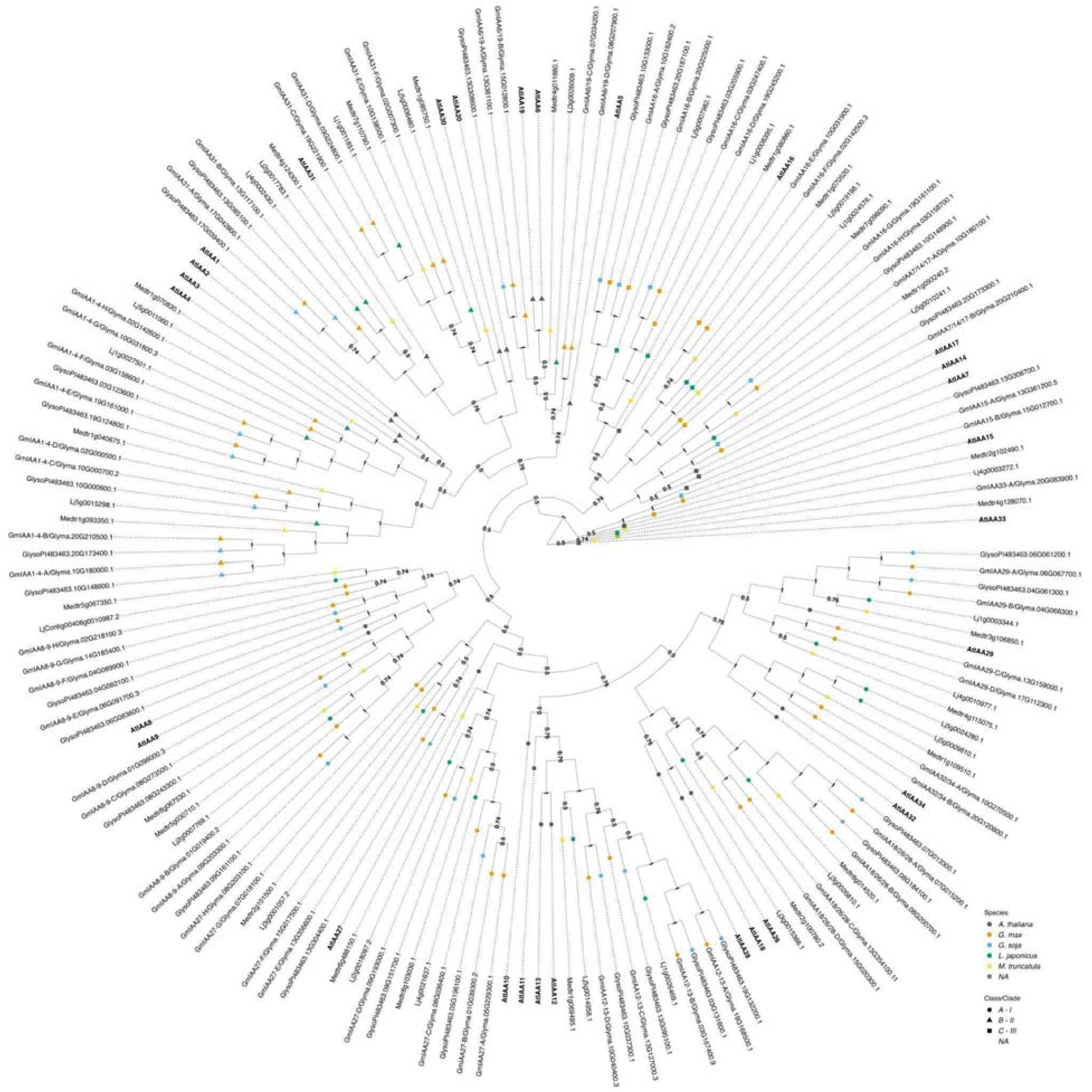
The evolutionary relationships between *G. max* Aux/IAA proteins, *A. thaliana* and other legume species orthologs. The historical relationship was inferred using the Bayesian Inference method (Ronquist *et al*., 2012). The optimal tree is drawn according to the posterior probability of the evolutionary distances. The posterior probability of each node is labeled. Each tip is colored according to species, with *A. thaliana* in black, *G. max* in orange, *G. soja* in light blue, *L. japonicus* in green, and *M. truncatula* in yellow. *A. thaliana* gene symbols are displayed in bold to better visualize assigned orthology. Aux/IAAs co-receptors are divided here into three classes/clades: Class A - I, represented as a circle; Class B - II, represented as a triangle, and class C - III, represented as a square. *G. max* were named according to both their orthology to *A. thaliana* followed by its gene ID. Nodes are labeled with their supporting probabilities in the center.

### Phylogenetic analysis of ARFs transcriptional factors and proposed orthology based on *A. thaliana* classification

The *ARF* gene family in *G. max* includes fifty-five members which are separated into three classes (Ulmasov *et al*., 1999; Tiwari *et al*. 2003; Finet *et al*., 2013). Class A ARFs are likely transcriptional activators and orthologous to AtARF5/6/7/8/19. The protein sequences of activator ARFs contain a Q-rich middle region that reinforces their activation properties (Appendix C). Class B ARFs are transcriptional repressors, and have a large clade containing AtARF1/2/3/4/9/11/12/13/14/15/18/20/21/22/23. Finally, class C, also composed of transcriptional repressors, contains AtARF10/16/17 and are nearest the root of the tree, likely as the split between class C ARFs and the remaining ARFs existed before the evolution of land plants (Figure 4) (Ulmasov *et al*., 1999; Finet *et al*., 2013; Flores-Sandoval *et al*., 2018; Mutte *et al*., 2018). As a result of *G. max* paleopolyploidy the majority of orthology groups have at least three *G. max* genes per corresponding *A. thaliana* ortholog. Additionally, all orthologs of ARF3, 8, 9, 10, 16, 7, and 19 *G. max* proteins have *G. soja* sisters, whereas the others lack at least one sister taxa in the wild relative. Interestingly, *M. truncatula* has ten orthologs of AtARF17, which is the outgroup in our analysis. This large group of orthologs is likely due to tandem duplications on *M. truncatula* chromosome 5. Similarly in *A. thaliana* ARF9, 12, 13, 14, 15, 20, 21, 22, and 23 likely resulted from tandem duplication events, and the legumes have many fewer genes in this clade, following the more typical pattern of whole genome duplication events.

**Figure 4.**
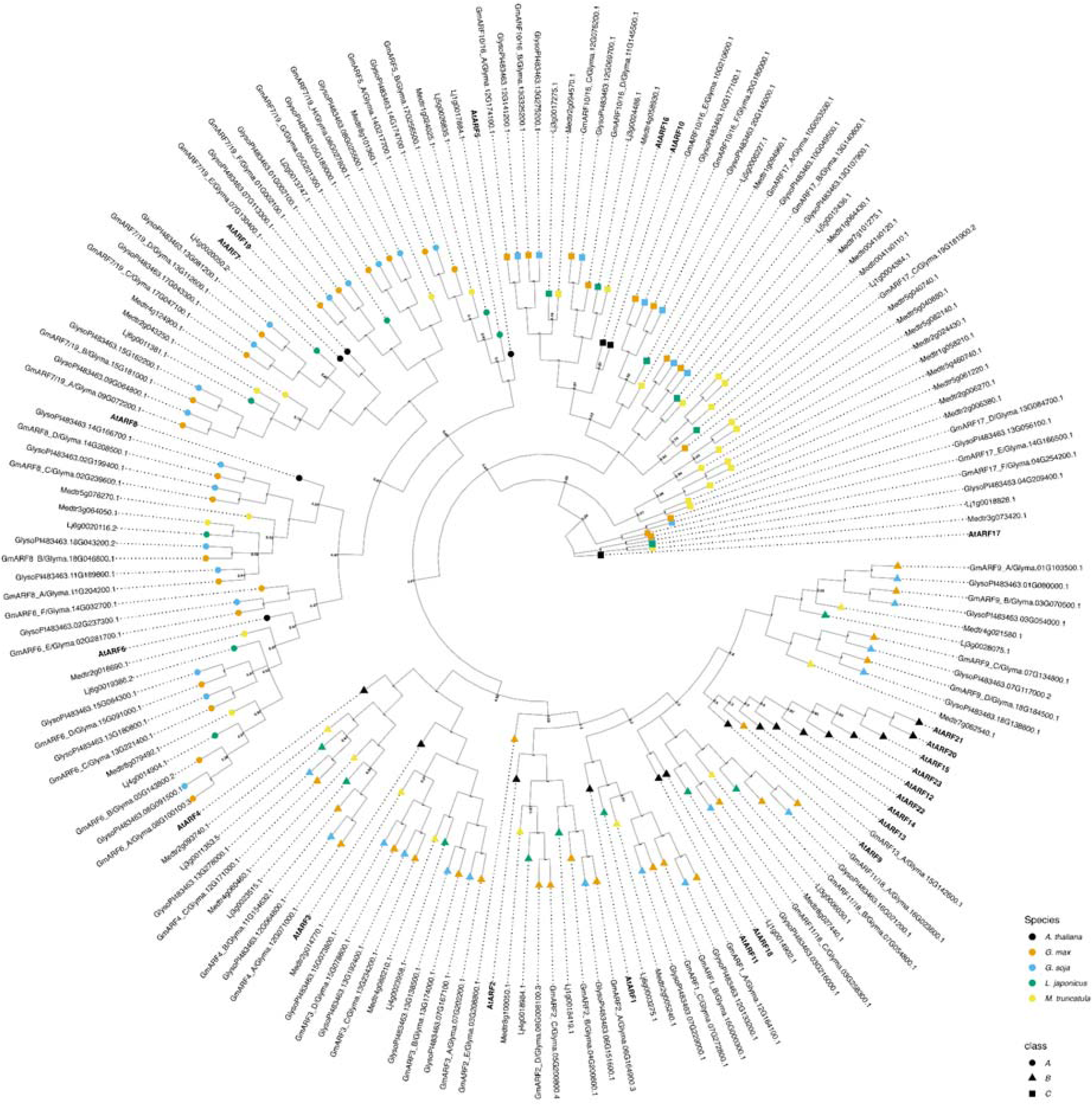
The evolutionary relationships between *G. max* ARFs proteins, *A. thaliana* and other legume species orthologs. The historical relationship was inferred using Bayesian Inference (Ronquist *et al*., 2012). The optimal tree is drawn according to the posterior probability of the evolutionary distances. The posterior probability of each node is labeled. Each tip is colored according to species, with *A. thaliana* in dark grey, *G. max* in orange, *G. soja* in light blue, *L. japonicus* in green, and *M. truncatula* in yellow. *A. thaliana* gene symbols are displayed in bold to better visualize assigned orthology. ARFs are divided here into three classes/clades (Ulmasov *et al*., 1999; Finet *et al*., 2013): Class A - II, represented as a circle; Class B - I, represented as a triangle, and class C - III, represented as a square. *G. max* were named according to both their orthology to *A. thaliana* followed by its gene ID.

## Discussion

Although the auxin signaling pathway is well studied and known to play an important role in plant growth, development, and architecture in A. thaliana, little is known about auxin’s roles in legumes (Li and Chen, 2023; Salehin et al., 2015) outside root and nodule development (Breakspear et al. 2014; Wang et al. 2015; Cai et al. 2017; Nadzieja et al. 2018; Schiessl et al. 2019; Wang et al. 2019; Gao et al. 2021; Rogato et al., 2021; Goto et al. 2022; Li and Chen, 2023) and shoot height/dwarfing. Auxin’s gene regulatory genes are composed of an extensive number of genes for each *TIR1/AFB*, *ARF*, and *Aux/IAA* families in both *A. thaliana* and *G. max*. These auxin genes work together in a tissue-dependent manner (Piya *et al*., 2014) conducting shoot development and rosette area (Vernoux *et al*., 2011; Salehin *et al*., 2015; Prigge *et al*., 2020), organ primordia, as well as cell fate (Rogato *et al*., 2021; Salehin *et al*., 2015; Parry *et al*., 2009) in plants. Therefore, auxin’s mechanism of regulation is a great candidate for plant breeding programs (Li and Chen, 2023). A lot has been achieved over the past decades in how the main components of the auxin signaling pathway (*TIR1/AFB* receptors, *Aux/IAA*s repressors and *ARF*s transcription factors) work together in conducting many aspects of plant growth and development (Vernoux *et al*., 2011; Salehin *et al*., 2015; Rogato *et al*., 2021). However, what novel complexity and/or functional redundancy contained in each of these gene families remains an open question.

We have identified 14 *TIR1/AFB and 4 COI1-like F-box*, 55 *ARF*s, and 61 *Aux/IAA*s members of *G. max* auxin signaling gene families based on their similarity/evolutionary history to *A. thaliana* genes. The evolutionary history of the TIR1/AFB proteins exhibited 5 clades in for the *G. max* genes which were clustered equivalently to results reported for most *A. thaliana*, *L. japonicus* and *M. truncatula* previous literature (Dharmasiri *et al*., 2005; Parry *et al*., 2009; Shen *et al*., 2015; Hamm *et al*., 2019; Rogato *et al*., 2021). To our knowledge, phylogenetic analysis of these TIR1/AFB auxin receptors found in the *G. max* genome have not yet been explored. We identified 61 *Aux/IAA* genes in *G. max*, which largely followed the clade structure of previous analyses (Remington *et al*., 2004; Liu *et al*., 2021; Ali *et al*., 2022). Similarly, we identified 55 *ARFs* in the *G. max* genome which fell into orthology groups largely as expected based on previous analyses (Le *et al*., 2016). Our orthology naming convention, which has not to our knowledge been established before for soybean, facilitated comparative evolutionary analysis of these families with that of *A. thaliana*.

### *TIR1/AFB* receptor genes important in *G. max* aerial architecture

Auxin transcriptional responses are governed by the degradation of *Aux/IAAs* in an auxin-dependent manner through interaction with *SCF^TIR1/AFB^* ubiquitin ligases and proteasomal degradation that releases class A activator *ARFs* from *Aux/IAAs* repression (Ramos *et al*., 2001; Zenser *et al*., 2001; Gray *et al*., 2001; Chapman and Estelle, 2009). Subsequently, different *TIR1/AFB-Aux/IAA-ARF* modules have been shown to regulate plant growth and development in a tissue-specific fashion in *A. thaliana* (Tatematsu *et al*., 2004; Vernoux *et al*., 2011; Piya *et al*., 2014; Krogan and Berleth, 2015; Galvan-Ampudia *et al*., 2020). For instance, whereas the control of gene expression in response to auxin in *A. thaliana* has shown to be governed by *ARF7* and *Aux/IAA3* heterodimerization in root development, in hypocotyl elongation the same *ARF7* preferentially dimerizes with *Aux/IAA19* (Tatematsu *et al*., 2004; Wilmoth *et al*., 2005). Analogously, in our PCA analysis, the proximity of *ARFs* and *TIR1/AFBs* to the origin (Figure 1) suggests their involvement in diverse tissue processes. Conversely, the *Aux/IAAs* exhibited greater dispersion, implying greater tissue specificity, and possibly engender distinct auxin responses in different tissues. Comparing expression levels within the *A. thaliana TIR1/AFB family*, *TIR1*, *AFB1*, *AFB2* and *AFB3* are all shown to be expressed in *A. thaliana* rosette leaves and meristematic regions, however expression of *AFB2/3* were the highest observed (Dharmasiri *et al*., 2005; Vernoux *et al*., 2011; Prigge *et al*., 2020). Additionally, AFB4 levels were nearly negligibly whereas AFB5 is involved in Aux/IAAs turnover in inflorescence meristems (Vernoux *et al*., 2011). We observed a similar pattern in *G. max* receptors with *GmTIR1/AFB1, GmAFB2/3* and *GmAFB4/5* orthologs highly expressed in meristematic regions and leaves (Figure S5). Although, we observe here that these genes are highly expressed in soybean shoots, we predict that GmTIR1/AFB1 orthologs primarily govern Aux/IAAs turnover in hypocotyl, cotyledon and leaves. In contrast *GmAFB2/3* orthologs appear to exhibit greater specificity within meristematic tissues (Figure 1). Hence, these GmAFB2/3 receptors may be important in regulating auxin-induced degradation of Aux/IAAs in tissues important for aerial plant architecture in *G. max* as they are in *A. thaliana*.

### Aux/IAAs repressors important in *G. max* aerial architecture

Interestingly, we observed several orthologs of *AtIAA16* that are highly and specifically expressed in hypocotyl, leaf, cotyledon, axillary and shoot apical meristems in soybean (Figure 1). Korasick *et al*. (2014) demonstrated that overexpressing AtIAA16 stunts vegetative growth, which can then be rescued by knocking out a binding face of the *IAA16* PB1 domain. Rinaldi *et al*. (2012) noted a dominant trait in *Atiaa16* gain-of-function mutants, which led to limited vegetative growth in adult plants. Based on Korasick’s *et al*. (2014) and Rinaldi *et al*. (2012) work, the high expression of *AtIAA16* orthologs in hypocotyl, leaf, cotyledon, axillary and shoot apical meristems in soybean may restrain apical growth. In addition, *AtIAA16* is predicted to interact with *AtARF8*, which is also closely related to *ARF6* and therefore likely share similar interaction patterns (Piya *et al*., 2014). Orthologs of AtARF6/8 were expressed highly in canopy cover associated soybean tissues mentioned above, underscoring the central role played by At*IAA16* orthologs in auxin signaling and the associated phenotypes (Figure 1). Similarly, *IAA7/14/17* are closely related to *IAA16*, and potentially interacts with activator *ARFs* (*ARF8/9*) (Piya *et al*., 2014; Korasick *et al*., 2014), and predicted to be an important asset in hypocotyl development in soybean (Figure 1). Likewise, *IAA17* was predicted to interact with *ARF1* and play a part in hypocotyl development in *A. thaliana* (Piya *et al*., 2014), which supports our findings that *IAA7/14/17* orthologs may be involved in hypocotyl development in soybean.

Currently, *Atiaa28* gain-of-function mutants show a strong phenotype for reducing apical dominance and plant size in arabidopsis (Rogg *et al*., 2001). This would be a noteworthy ortholog to study in other plants. Likewise, gain-of-function mutants of *Gmiaa27* are known to influence apical dominance and branching in soybean plants (Su *et al*., 2022). While we did not detect orthologs of *GmIAA27* in our PCA, its high expression in meristems is evident (Figure S5). Additionally, *SiIAA19* has been linked to multiple auxin signaling processes, such as apical dominance (Sun *et al.,* 2013). It is possible that orthologs of these genes may serve as interesting targets as well. PAP1 or IAA26 in arabidopsis has also been linked to apical dominance due to loss of the trait after RNA silencing (Padmanabhan *et al*., 2005). Likewise, we saw in our heatmap (Figure S5) that IAA19/26/28 are highly expressed in leaf, and shoot apical meristem in soybean. Interestingly, all of these orthologs are phylogenetically-related, reaffirming what was discussed by Piya *et al*. (2014) that closely related proteins are prone to display similar modes of action.

Most ARF family members can form complexes with most Aux/IAA family members interchangeably (Piya *et al*., 2014). Negative feedback in the greater auxin signaling network also prevents quantification of molecular function in planta. Further investigation is needed to clarify the mechanism of specific ARF and AUX/IAA family members on transcriptional dynamics. Due to these confounding factors, pinpointing many Aux/IAA orthologs will be crucial for finding some that can be harnessed for rational engineering of plant growth.

### ARFs transcription factors important in *G. max* aerial architecture

*AtARF2, AtARF8,* and *AtARF9* orthologs stand out in our analyses as being highly and specifically expressed in RCC tissues in *G. max*. As stated above *GmARF2_A.2/B.1/A.3/C.4/D.2* and *GmARF9_B.2* were associated with meristematic tissues. *GmARF9_B.1, ARF2_C.1*, and *ARF8_C.1* are strongly associated with hypocotyl, leaf and cotyledon tissues. Described below, many of our findings are corroborated by existing literature by means of phenotypic and genomic analyses in *Arabidopsis, G. max*, and other related species. However, our analysis suggests that seemingly redundant *ARF* paralogs may have also evolved unique roles in *G. max.* Additionally, despite extensive genetic analyses of the ARF protein family (Okushima et al., 2005b), the delineation of *ARF* functionality in the SAM has been primarily limited to *ARF5*, while the involvement of other *ARF* genes is mostly supported by indirect evidence (Hardtke 2004; Mallory 2005). Our analysis provides more evidence in corroboration of such conclusions.

AtARF2, a class B ARF, is thought to serve as a negative regulator of cell proliferation and enlargement. In seedlings, AtARF2 serves as a bridge between ethylene and auxin signaling, playing a key role in apical hook formation (Li et al., 2004), backing our data suggesting the association of *GmARF2_C.1* with hypocotyl tissues. Furthermore, *ARF2 orthologs* serve as a regulator of leaf senescence in both *G. max* and *Arabidopsis* (Lim et al., 2010; La et al., 2022). Mutations in *AtARF2* result in delayed leaf senescence by reducing the repression of auxin signaling and increasing auxin sensitivity (Lim et al. 2010). Mutations in *AtARF2*, akin to loss-of-function achieved through gene silencing, led to elongated hypocotyls, darker green rosette leaves, and enlarged cotyledons, but did not impact global expression of auxin regulated genes in Arabidopsis seedlings (Okushima et al., 2005a). *GmARF2, represented here by its variants GmARF2_A.2/A.3,* expression was upregulated in shaded *G. max* plants, contributing to leaf enlargement inhibition (Wu et al., 2017). *AtARF2* has also been cited as playing a role in the SAM. For example, in Arabidopsis, SAM cells are maintained during embryogenesis by down-regulating ARF2 activity (Roodbarkelari et al., 2015). This corresponds with the tissue association of GmARF2_A.2/B.1/A.3/C.4/D.2 in our PC analysis *AtARF8* orthologs were the only class A ARFs that were strongly associated with any of the tissues in our analysis. Regulated by photoreceptors, CRY1 and phyB in *Arabidopsis*, *AtARF6* and *AtARF8* in turn are associated with regulation of hypocotyl elongation under blue and red light. *AtARF8/ARF6* double null mutants also have reduced responses to environmental conditions. Far red light and elevated temperature exposure stunt hypocotyl elongation (Mao et al., 2019). AtARF8, in conjunction with AtARF6, indirectly mediates the expression of a key brassinosteroid (BR) biosynthetic enzyme in *Arabidopsis*, which ultimately directs proximodistal cell expansion (Xiong et al., 2021). Leaf shape is primarily determined by proximodistal growth. BRs also promote cell wall loosening which has been shown in simulations to lead to cell and organ growth, and thus modulate leaf roundness (Xiong et al., 2021). AtARF8 operates redundantly with AtARF6 to repress phloem proliferation and induce cambium senescence during the xylem expansion phase in the hypocotyl by interacting with DELLA proteins from the gibberellin signaling pathway. AtARF8 and AtARF6 also play an essential role in cambium establishment and maintenance (Ben-Targem et al., 2021). In *M. truncatula*, the *ARF8* ortholog exhibits slightly elevated expression in the petiole and stem, but not the leaf (Liu et al., 2021). Similarly, our analysis found a slight association of *GmARF8_C.1* with hypocotyl, leaf and cotyledon tissues. We also discovered a stronger association of GmARF8_C.3 with the SAM, along with GmARF6_C.2 and GmARF8_A.1 with the meristem and leaf (Figure S6). AtARF6 and AtARF8 are typically cited together due to their redundant expression domain and functionality, which is observed to some extent in our analysis of *G. max*.

One ortholog of *AtARF9*, *GmARF9_B.1*, another class B repressor, stood out in our analysis as potentially playing a role in shoot architecture. *GmARF9,* here *GmARF9_C.1* variant, is associated with promoting first pod height (Jiang *et al*., 2018). In *M. truncatula*, *MtARF9* has elevated expression levels in the leaves, shoots, and petioles among other tissues in the roots and seeds (Liu et al., 2021). There is otherwise a notable lack of literature that draws any meaningful connection between *ARF9* and shoot architecture. Nonetheless, our PC analysis suggests that several of the *GmARF9* paralogs may serve distinct roles in the SAM, hypocotyl, leaf, and cotyledon tissues. However, Vernoux et al. found that *AtARF9* has a fairly weak homogenous expression pattern in the *Arabidopsis* SAM, postulating that *ARF9* likely does not play a dominant role in that tissue (Vernoux et al., 2011). This discrepancy between the literature and our results could be explained by divergent roles of gene paralogs. Our analysis may also point to previously unknown functions of *GmARF9_B.1* and its role in regulating shoot architecture.

*ARF2,8,9* paralogs in *G. max* may serve as key targets in future studies exploring the developmental regulation of RCC. Further investigation is needed to clarify the roles of specific ARF-mediated transcriptional dynamics which is further confounded by the complex network of interactions with the large family of Aux/IAA proteins which play a key role in modulating unique transcriptional responses.

In conclusion, the findings presented in this study pinpoint potential auxin candidate genes that hold promise for addressing RCC development in soybean. While we are enthusiastic about these results, we recognize the constraints of our analysis, primarily stemming from the limited available data. To advance our comprehension and the future application of this research, it is essential to conduct further investigations on these genes and their spatiotemporal expression patterns. Additionally, we plan to employ a yeast heterologous system to functionally characterize RCC-related TIR1/AFB-Aux/IAA-ARFs modules. This approach will enable us to finely tune auxin responses and gain deeper insights into its intricate interaction network. Subsequently, we can apply this newfound knowledge to in planta studies, facilitating the correlation of phenotypes with this complex network.

## Methods

### Sequence collection

Seven AFB, twenty-three ARF, and twenty-nine aux/IAA’s *A. thaliana* amino acid sequence identities from Hamm *et al*. (2019) were used in sequence retrieval through the BSgenome.Athaliana.TAIR.TAIR9 and r1001genomes R packages (Team TBD, 2014; Hamm *et al*., 2019). The peptide sequences were used in the initial Basic Local Alignment Search Tool (BLAST) against the *Glycine max* (assembly Wm82.a4.v1), *Glycine soja* (assembly Gsoja_v1_1), *Medicago truncatula* (assembly Mtruncatula_Mt4_0v1), *Lotus japonicus* (assembly Ljaponicus_Lj1_0v1), as well as *Arabidopsis thaliana* (assembly Athaliana_Araport11) genomes database in Phytozome V13 (Goodstein *et al*., 2012). Peptide sequences with an E-value less than or equal to 1E–50 were used for analysis. ARF and Aux/IAA sequences were manually separated in some cases according to their length, and the presence of a “QVVGWPPv/i” canonical *Aux/IAA* degron, or B3 ARF DNA binding domain. Retrieved peptides in ARFs search containing less than 400 base pairs (bp) or containing auxin canonical degron was removed from fasta file.

### Sequence alignment

Amino acid (AA) sequence alignment was performed using the function AlignSeqs from DECIPHER 2.24.0 R package (Wright, 2015) following default parameters. The alignment was built according to a similarity tree based on pairwise distinction of shared AA sequences. Two iterations followed the three built in which sequences were re-aligned to the three until convergence was reached. Finally, there was a refinement step in which portions of the alignment were re-aligned to the remnant of the alignment where two alignments were generated and the one that reached convergence with the best sum-of-pairs score was kept (Wright, 2015). Ultimately, we accounted for low information portions of the alignment due to highly-variable and/or gap regions by applying the MaskAlignment function in order to remove those regions. All settings, except for windowSize equals to 6 in MaskAlignment, followed default recommendations.

### Phylogenetic analysis

A nexus file was written from masked sequences using write.nexus.data function from ape’s R package version 5.6-2 (Maddison *et al*., 1997) for building the phylogeny trees. We then built phylogenies on MrBayes v3.2 software (Ronquist *et al*., 2012) based on the provided peptide sequences in the nexus file of the homologous proteins of AtAFB, AtARF, and AtAux/IAA family members for *G. max*, *G. soja*, *M. truncatula*, and *L. japonicus*. We defined AtCOI1, AtARF17, and AtIAA33 as outgroups to build AFB, ARF, and IAA Bayesian Markov chain Monte Carlo (MCMC) phylogenies, respectively. The likelihood model was defined using lset command with nucleotide substitution model set to protein and rates took into account a gamma distribution. The prior probability for the evolutionary model was defined using a fixed protein model (Jones model) and the proposal probability set to zero. The posterior probability of the phylogenetic trees were calculated based on the MCMC parameters: for TIR1/AFBs we used a critical value for topological convergence diagnostic of 0.01, 6 chains, and the markov chain was sampled at every 100 cycles, additionally 0.25 of samples were discarded when convergence diagnostic was calculated.For the ARFs, the sample frequency was increased to 10000, number of runs set to 2, minimum frequency partition set to 0.05 and number of chains equal to 8. Finally, we followed the same parameters for Aux/IAAs, except that we increased the number of chains to 12. Ultimately, phylogenies visualization and annotation were drawn using ggtree and ggplot2 R packages (Wickham *et al*., 2016). Ortholog analysis based on the resulting phylogenies was used to assign *A. thaliana* ortholog names for each *G. max* gene. Table 1 provides a correspondence table of ortholog names to Wm82.a4.v1 gene IDs.

### Expression analysis data

Eleven soybean tissues RNA-seq raw data, project PRJNA241144, were downloaded from NCBI SRA repository and three other tissues RNA-seq data were downloaded from soybase. Soybean open flower (OF), inflorescence before and after meiosis (IBM and IAM), callus, hypocotyl, cotyledon, root tip, axillary meristem (AM), as well as shoot apical meristem at 6, 17, and 38 days (SAM6D, SAM17D, AND SAM38G) were obtained in NCBI, whereas root, young leaf and nodule tissues raw data in soybase. RNA-seq data was used for tissue-specific analysis using Galaxy *Salmon quant* tool. The raw gene expression counts were normalized to Transcripts Per Million (TPM). Normalization of the expression data to TPM allowed with ease the comparison of a gene expression between samples. The normalized data from Salmon quant tool was then manipulated in R using the pheatmap function from ggtree version 3.4.1 (Yu, 2022). The heatmap was built using median expression across all tissues equal or greater than 2, and normalized gene expression. Normalization was performed according to the following formula:

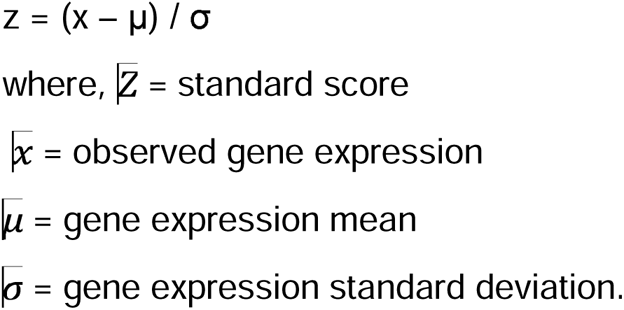

In order to further analyze and identify which genes contribute to rapid canopy cover related tissues we used the principal component analysis (PCA) unsupervised method. PCA of the gene expression were calculated using the *prcomp* function of the stats R package version 4.2.C (R Core Team, 2023). Parameters used for *prcomp* included *center* and *scale* set to *true*, meaning that the result is a correlation-based PCA. PCA retrieves loading segments through the linear combination of the gene expression values, thus reducing the dimensions of the data set. Additionally, PCA can be used to identify similarities and dissimilarities between genes and their contribution to each principal component and its respective loadings or eigenvectors. The data fed in the PCA calculations were all transcripts in which median expression for the fourteen tissues in questions were equal or greater than 2, resulting in a total of 133 transcripts well expressed. PCA presented in this study show the predicted genes for the seven tissues important in aerial growth, yet we show in supplemental data a PCA pertinent to the fourteen tissues. PCA plots were built using the *ggbiplot* function of the ggbiplot R package version 0.55 (Vu, 2011). Parameters used for *ggbiplot* included an ellipse set to *true* and *ellipse.prob* set to *70%* confidence interval. Ellipses were used as a visual representation of gene expression (data points) dispersion within each group in the PCA. Additionally, ellipses are drawn based on the covariance structure of gene expression for each group, meaning that size and orientation of the ellipses are determined by the covariance matrix. Finally, a table containing all gene information, along with data used in expression analysis was also built.

## Supporting information

https://github.com/PlantSynBioLab/SoyARC_manuscript_public/tree/bb41455d2204d228289bd31c90031190f5065753/Ready%20for%20Publication%20Files

## Data availability statement

We used RNA-seq raw data from project PRJNA241144 downloaded from NCBI SRA repository and from soybase. Additionally Scripts written for all data analysis can be found at https://github.com/PlantSynBioLab/SoyARC_manuscript_public/releases/tag/Auxin.

## Acknowledgments

Research in the Wright Plant Synthetic Biology Laboratory: the National Institute of General Medical Sciences of the National Institutes of Health under award number R35GM150856, the United States Department of Agriculture National Institute of Food and Agriculture (USDA-NIFA), Agriculture and Food Research Initiative (AFRI) Plant Breeding for Agricultural Production Grant No. 2022-67013-36293 and Hatch Project [VA-1021738]; Virginia Space Grant Consortium; Virginia Tech Institute for Critical Technologies and Sciences Junior Faculty Award; and Virginia Tech College of Agriculture and Life Sciences Strategic Plan Advancement 2021 Integrated Internal Competitive Grants through the Center for Advanced Innovation in Agriculture.

## Author Contributions

D. F. N. Planned and performed data analysis, and wrote the manuscript;

J. T. wrote some of the manuscript;

J. B. Planned some of initial data exploration, and wrote some of the manuscript;

B. B. Supervised some of the work and wrote some of the manuscript;

R. C. W. Supervised the work and wrote some of the manuscript.

All authors contributed to the interpretation of the data and revised and approved the final manuscript.

---

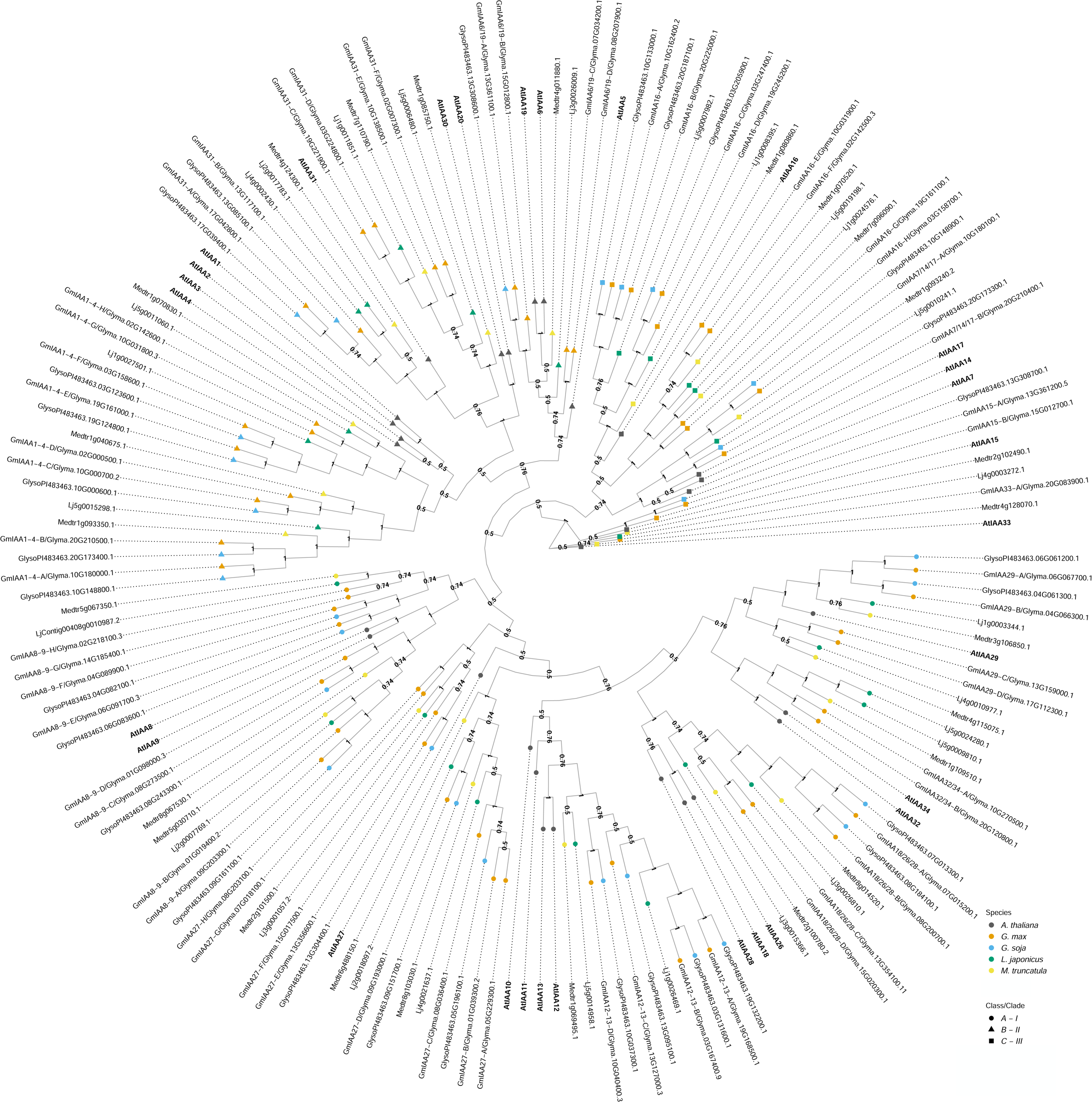

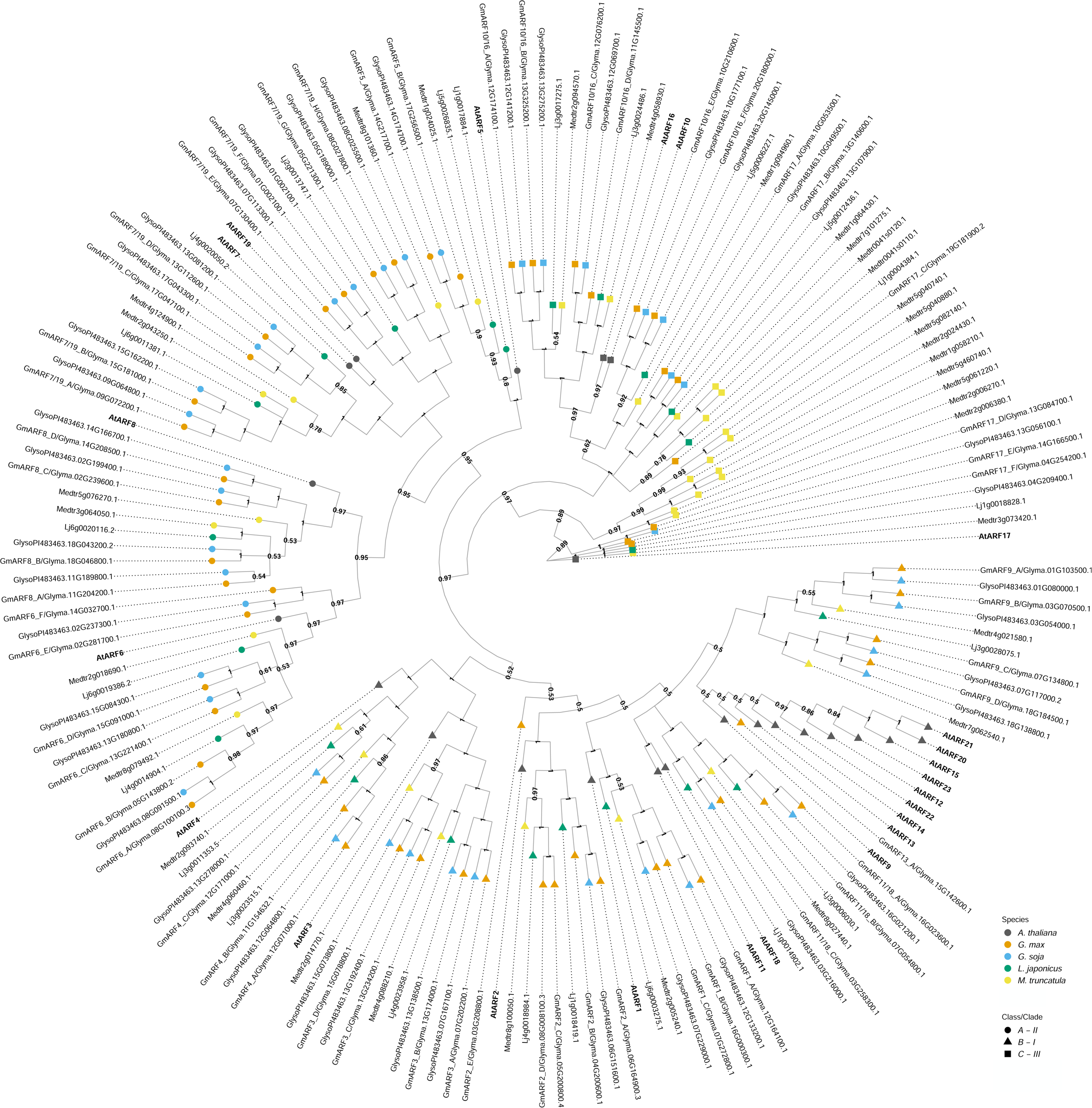

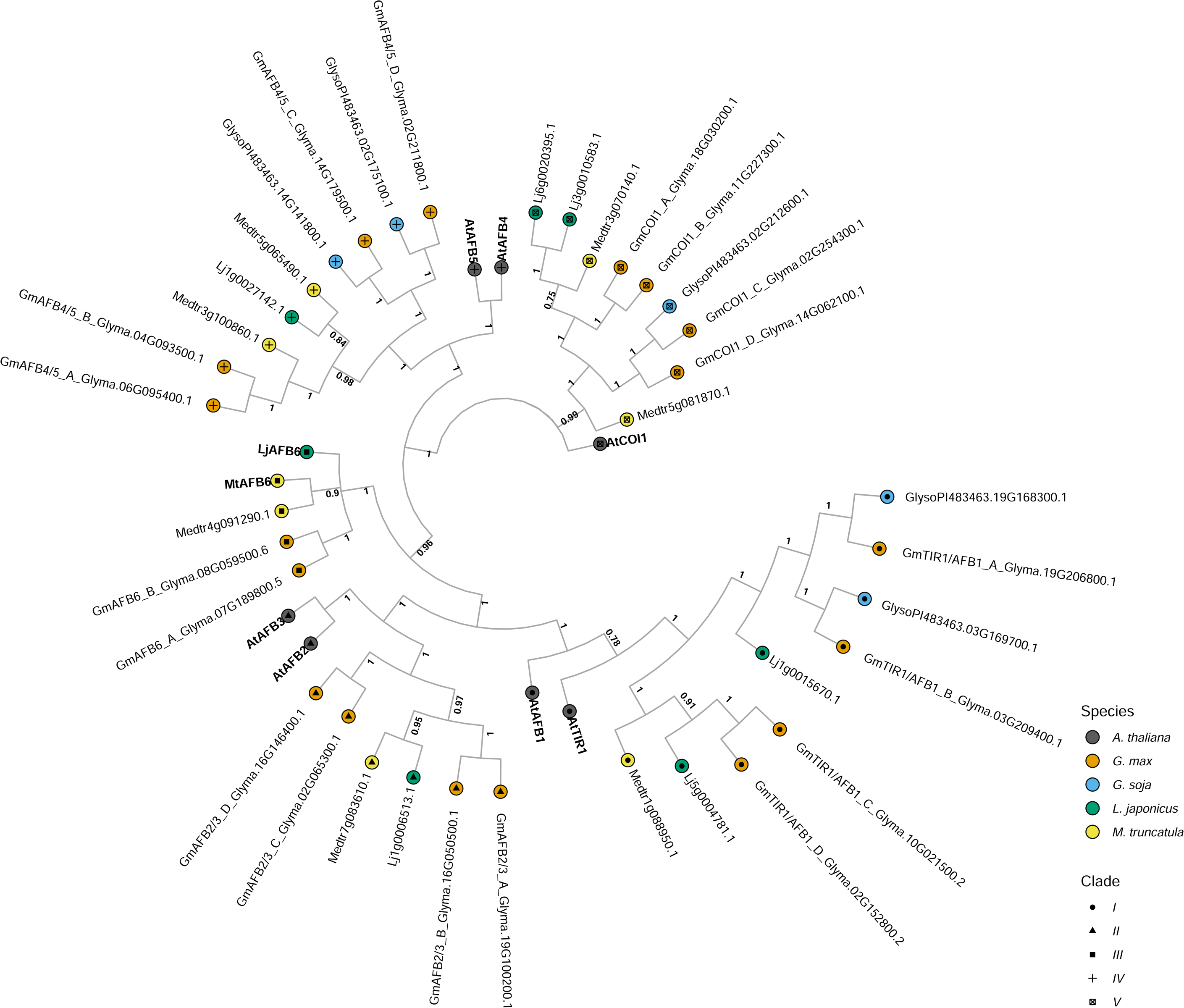

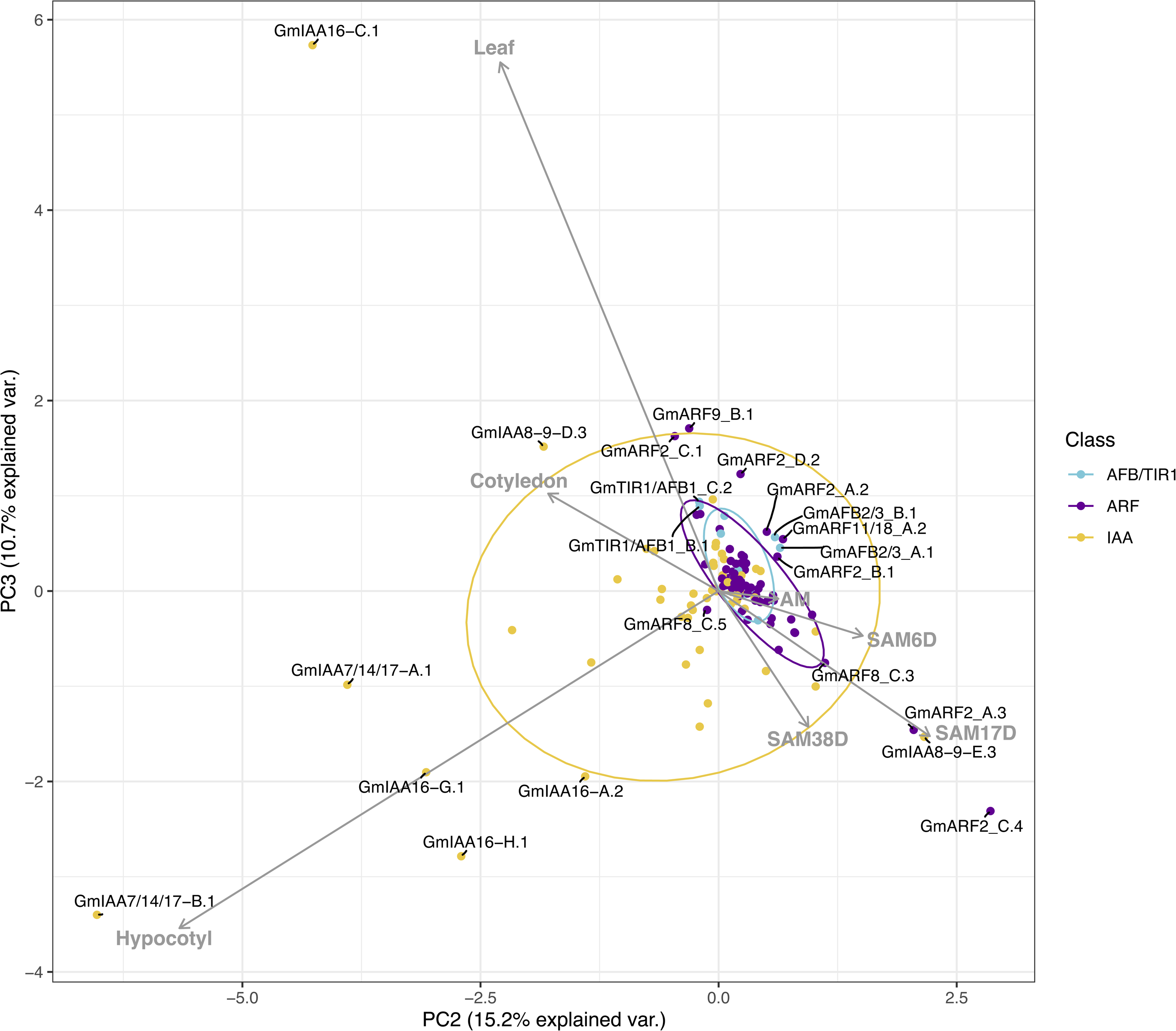

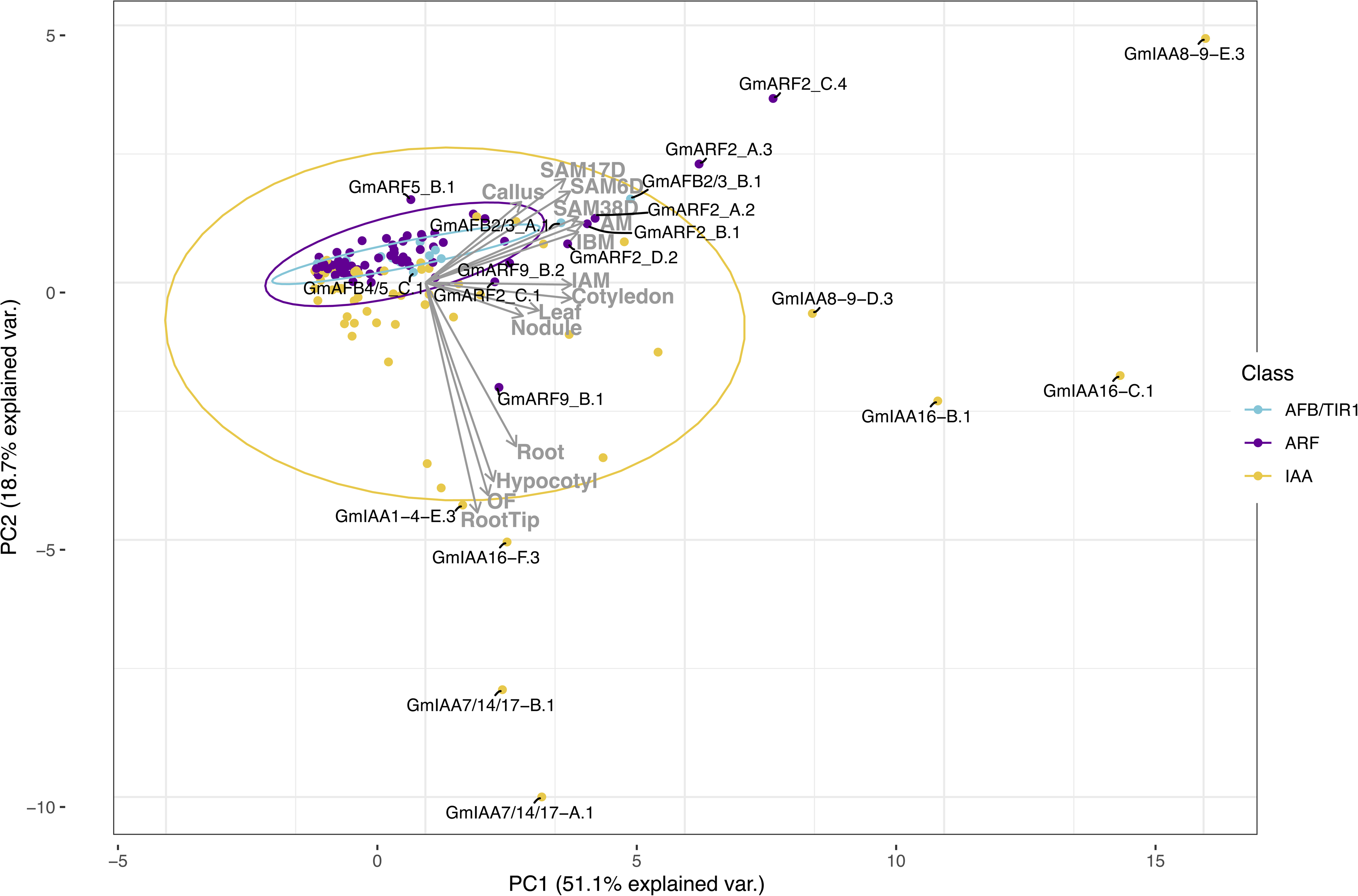

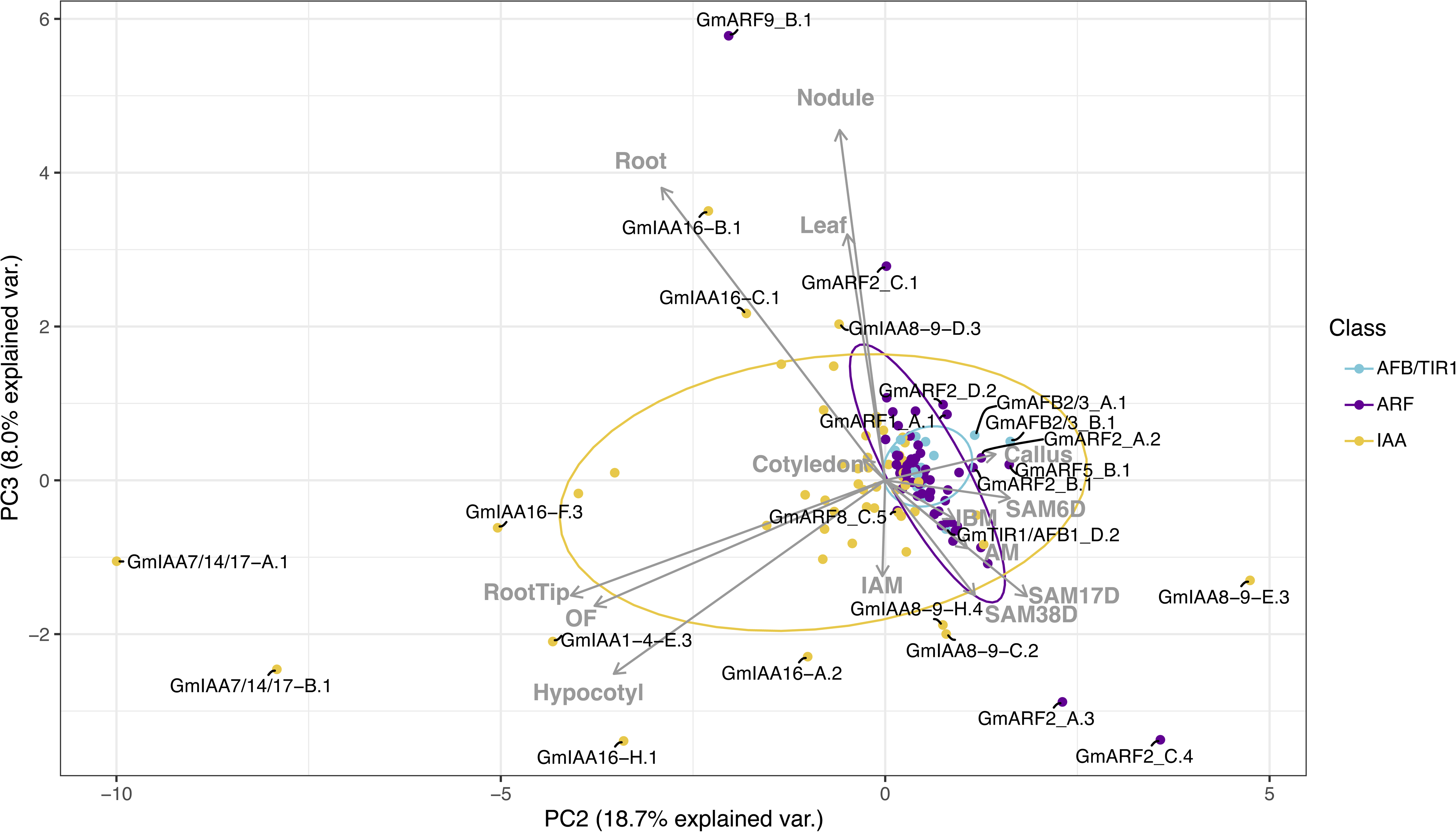

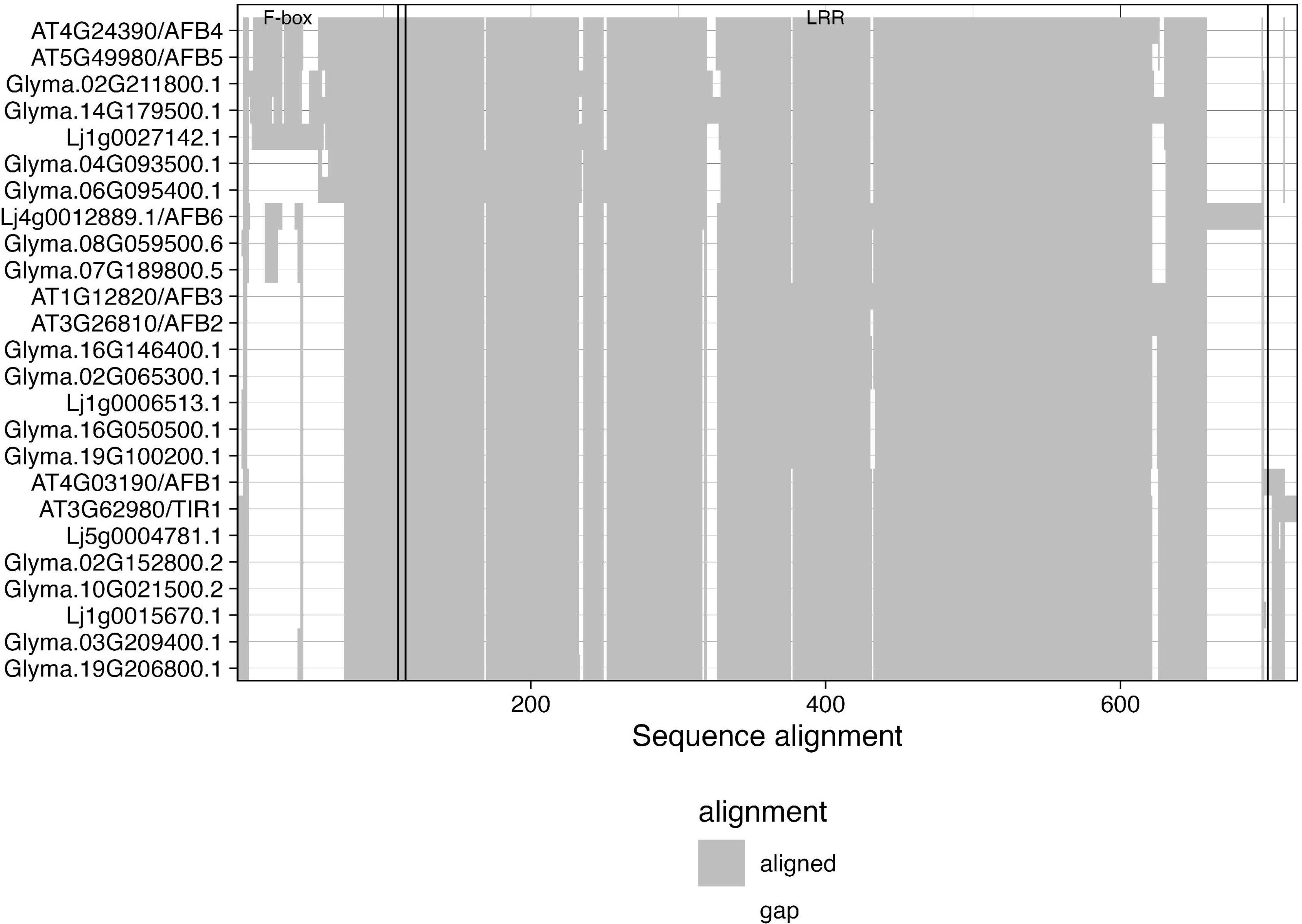

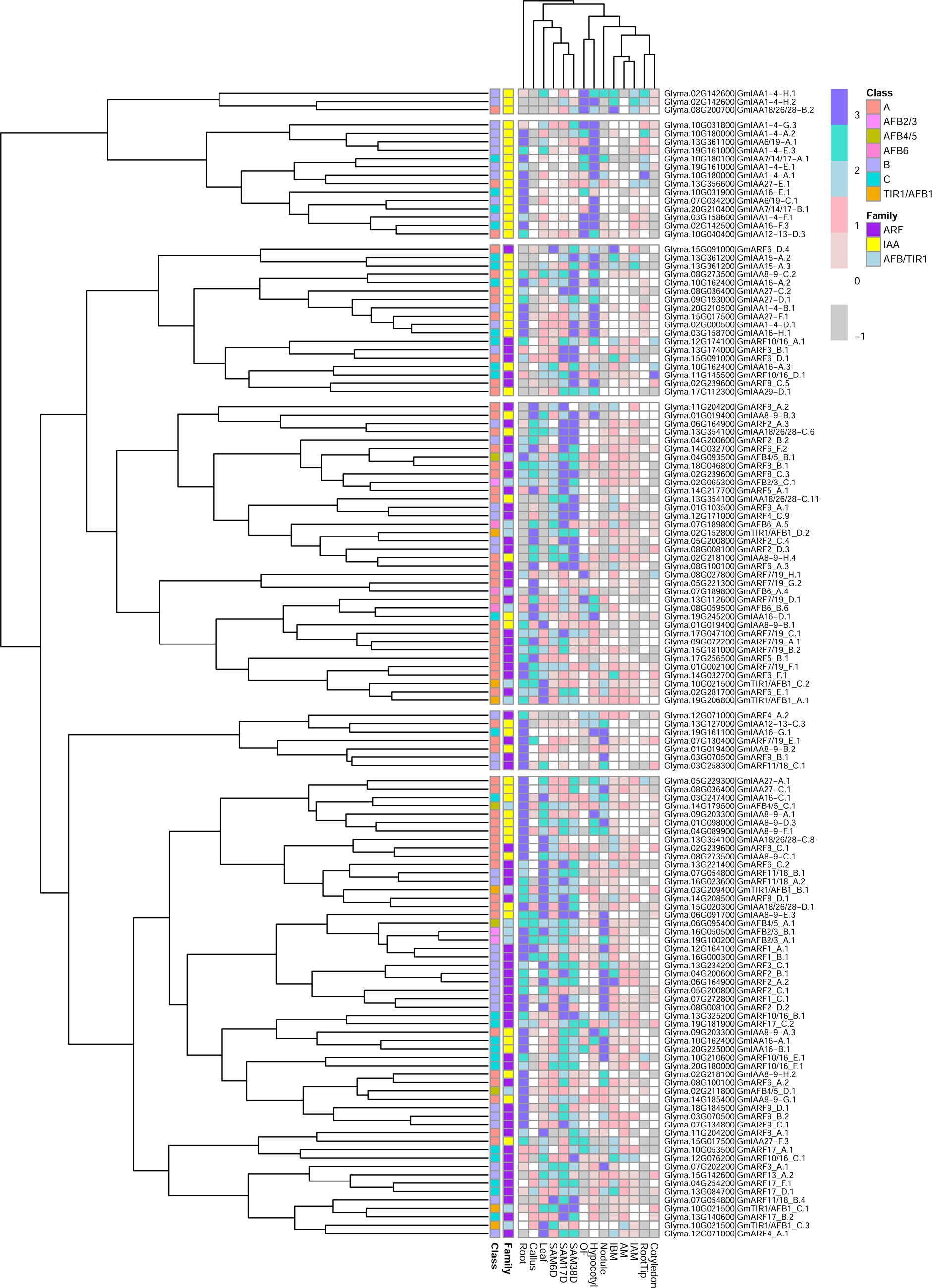

