## Supplementary material for "Identification of potential Auxin Response Candidate genes for soybean rapid canopy coverage through comparative evolution and expression analysis": https://github.com/PlantSynBioLab/SoyARC_manuscript_public/tree/bb41455d2204d228289bd31c90031190f5065753/Ready%20for%20Publication%20Files

Supplementary Figures

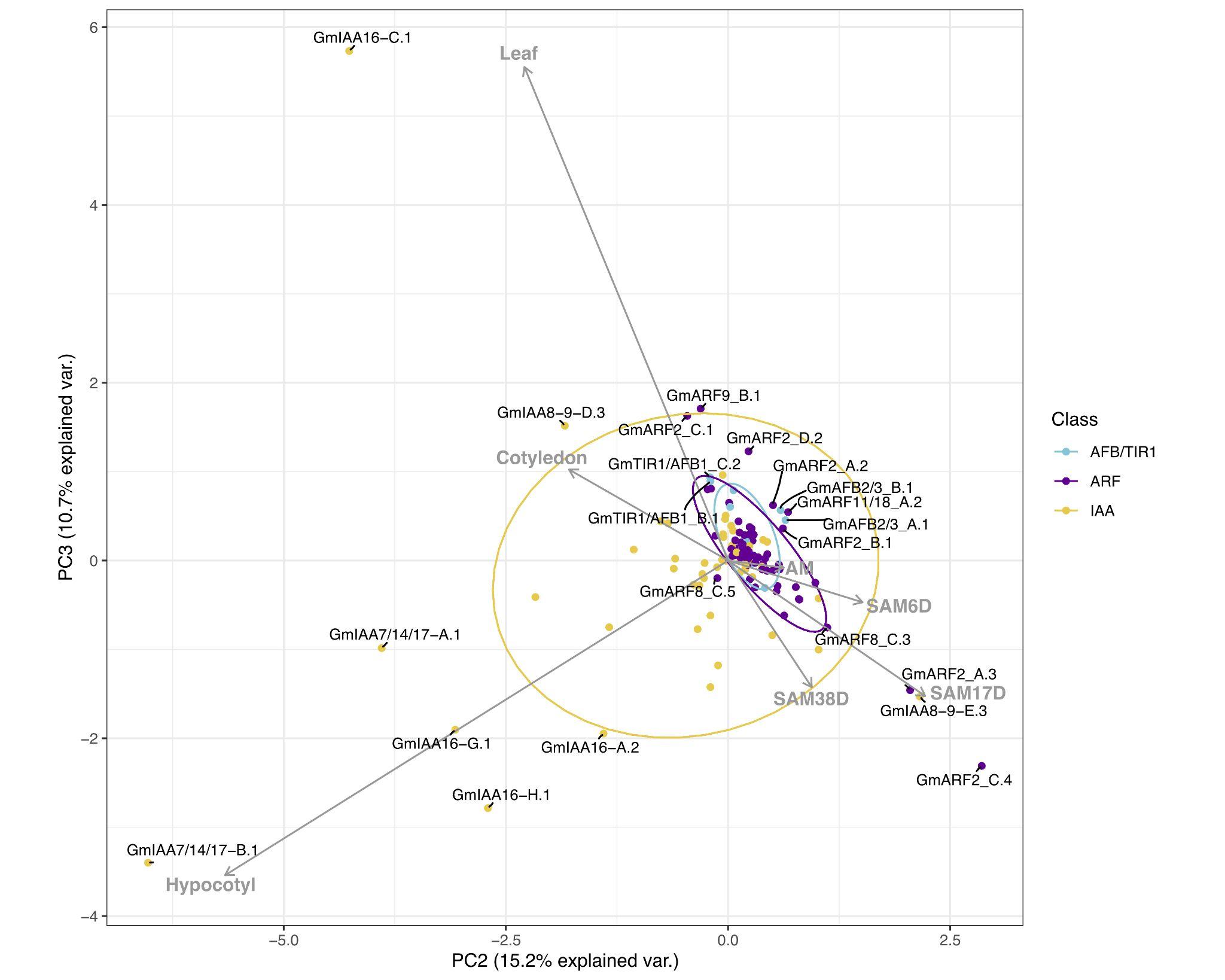

Figure S1. Correlation-based Principal Component Analysis (PCA): biplot of gene transcript expression and explanatory tissues involved in plant aerial architecture as eigenvectors (grey arrows), n = 7). Principal components 2 and 3 account for 25.9% of the total inertia. Ellipses are used here as a visual representation of dispersion of data points within each group (TIR1/AFB, ARF, and Aux/IAA (IAA)) with a 70% confidence interval. TIR1/AFB genes are colored cyan, ARF genes are colored purple, and Aux/IAA (IAA) genes are colored yellow. Some labels are connected to their respective points with hard lines. Genes clustering together inside the ellipses are hypothesized to have more pleiotropic effects on plant growth and development, whereas genes associated with a specific RCC tissue (genes that fall along an eigenvector, outside of the respectively colored ellipse) are hypothesized to have narrower effects and be more amenable to engineering RCC traits through gene editing.

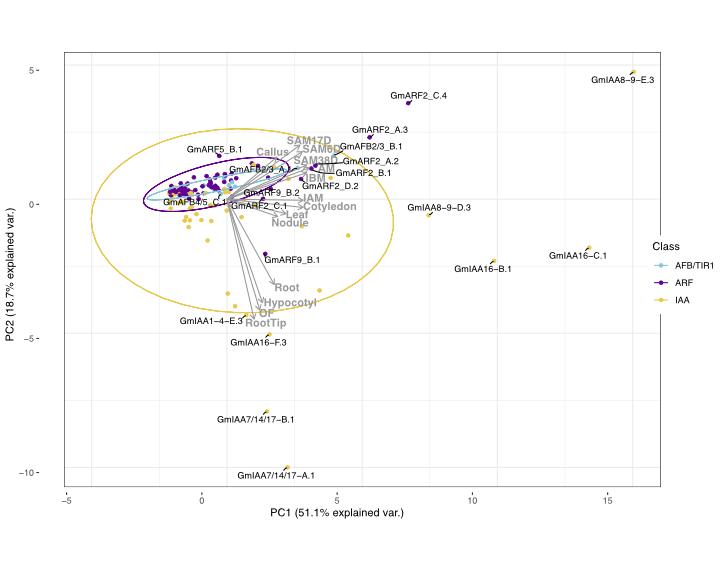
Figure S2. Correlation-based Principal Component Analysis (PCA): biplot of gene transcript expression and explanatory tissues involved in plant growth as eigenvectors (grey arrows), n = 14). Principal components 1 and 2 account for 69.8% of the total inertia.

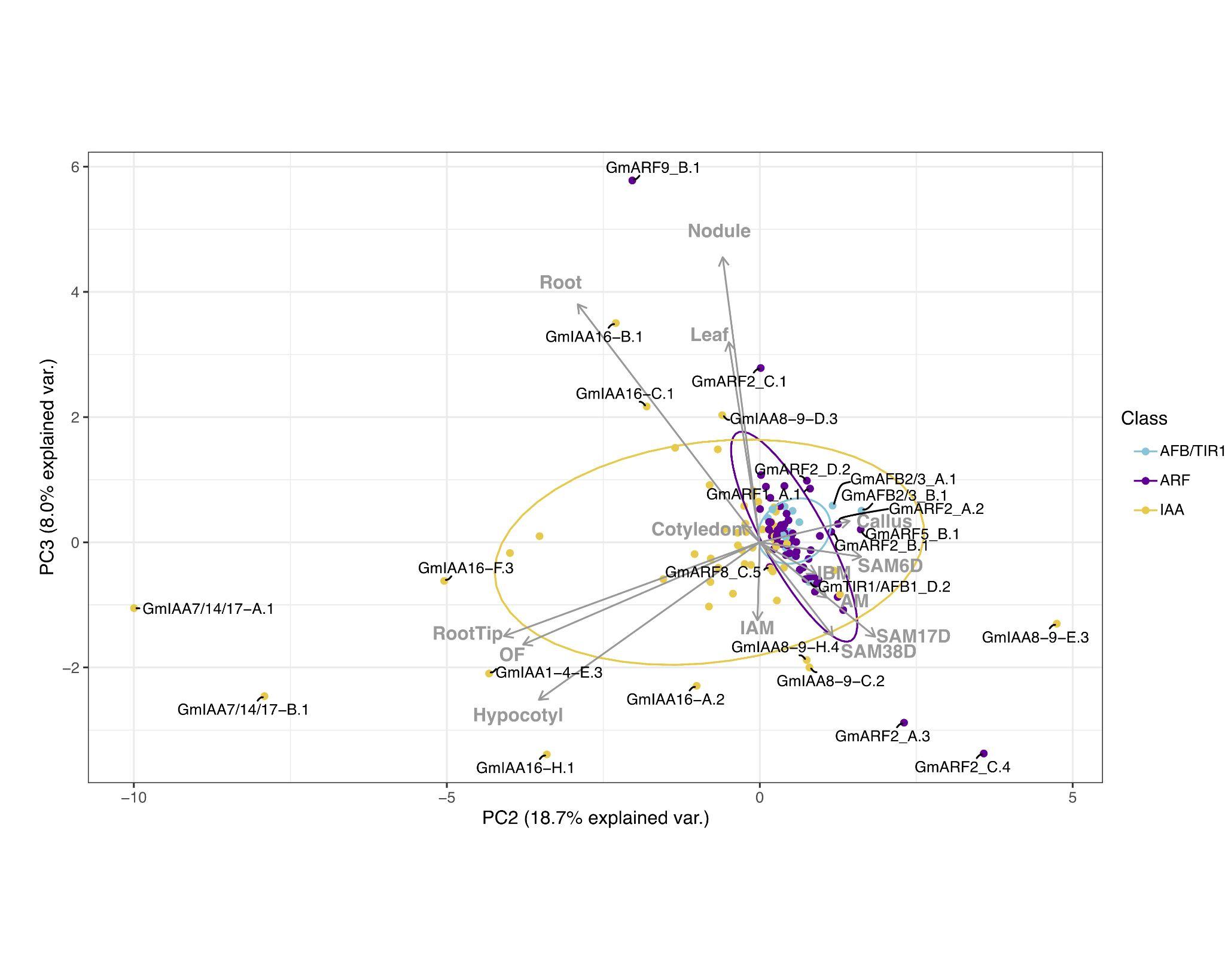

Figure S3. Correlation-based Principal Component Analysis (PCA): biplot of gene transcript expression and explanatory tissues involved in plant growth as eigenvectors (grey arrows), n = 14). Principal components 2 and 3 account for 26.7% of the total inertia.

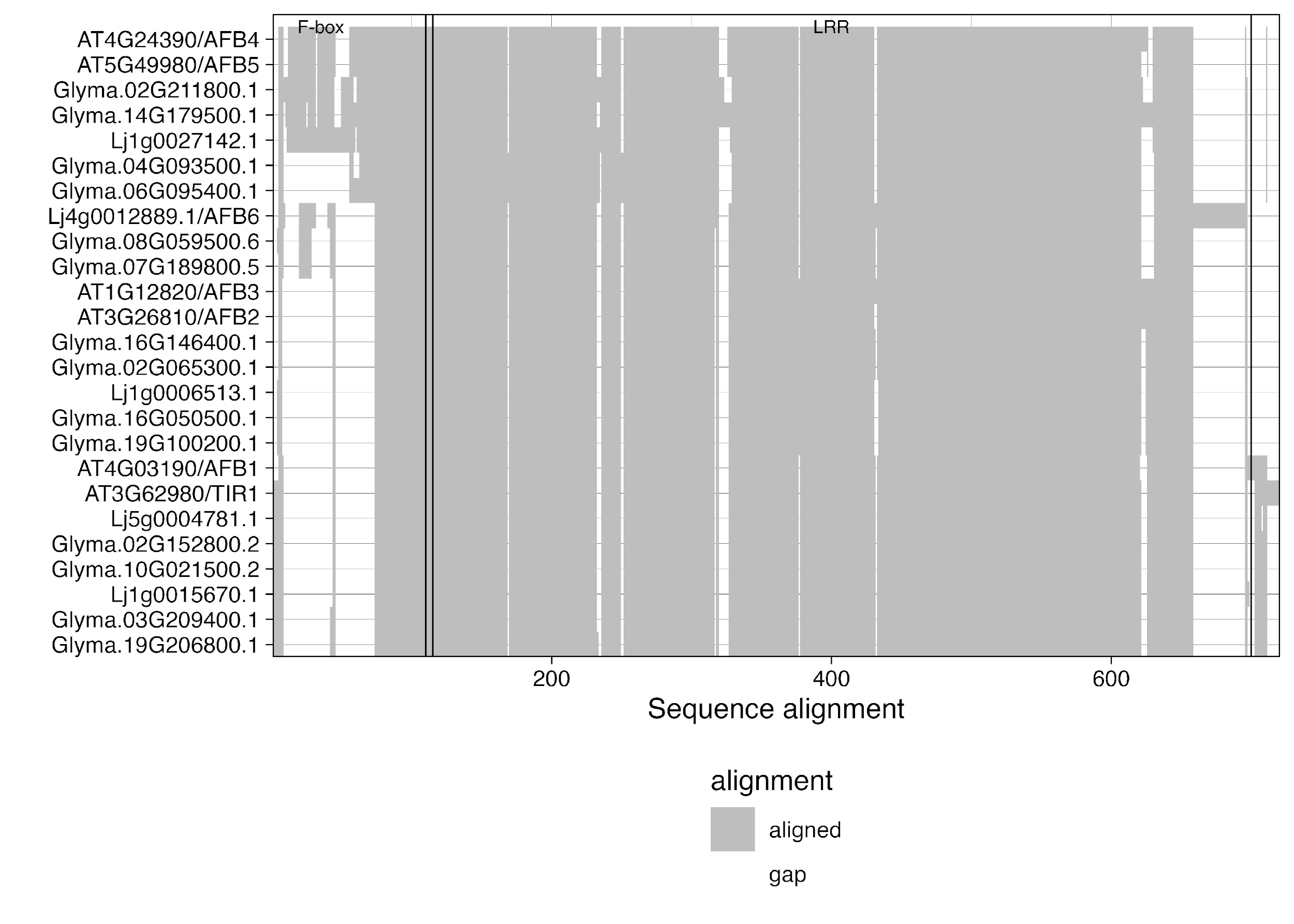

Figure S4. Critical functional domain similarity shared between *A. thaliana, L. japonicus* and *G. max* TIR1/AFB protein sequences. Conserved critical domains are depicted by the F-box label and its black line marking the end of it, which then follows a second black line demarking the start of the LRR region. Complete alignment showing conserved amino acids can be found in Appendix A. Protein sequences were aligned with DECIPHER, Wright (2015), in both cases.

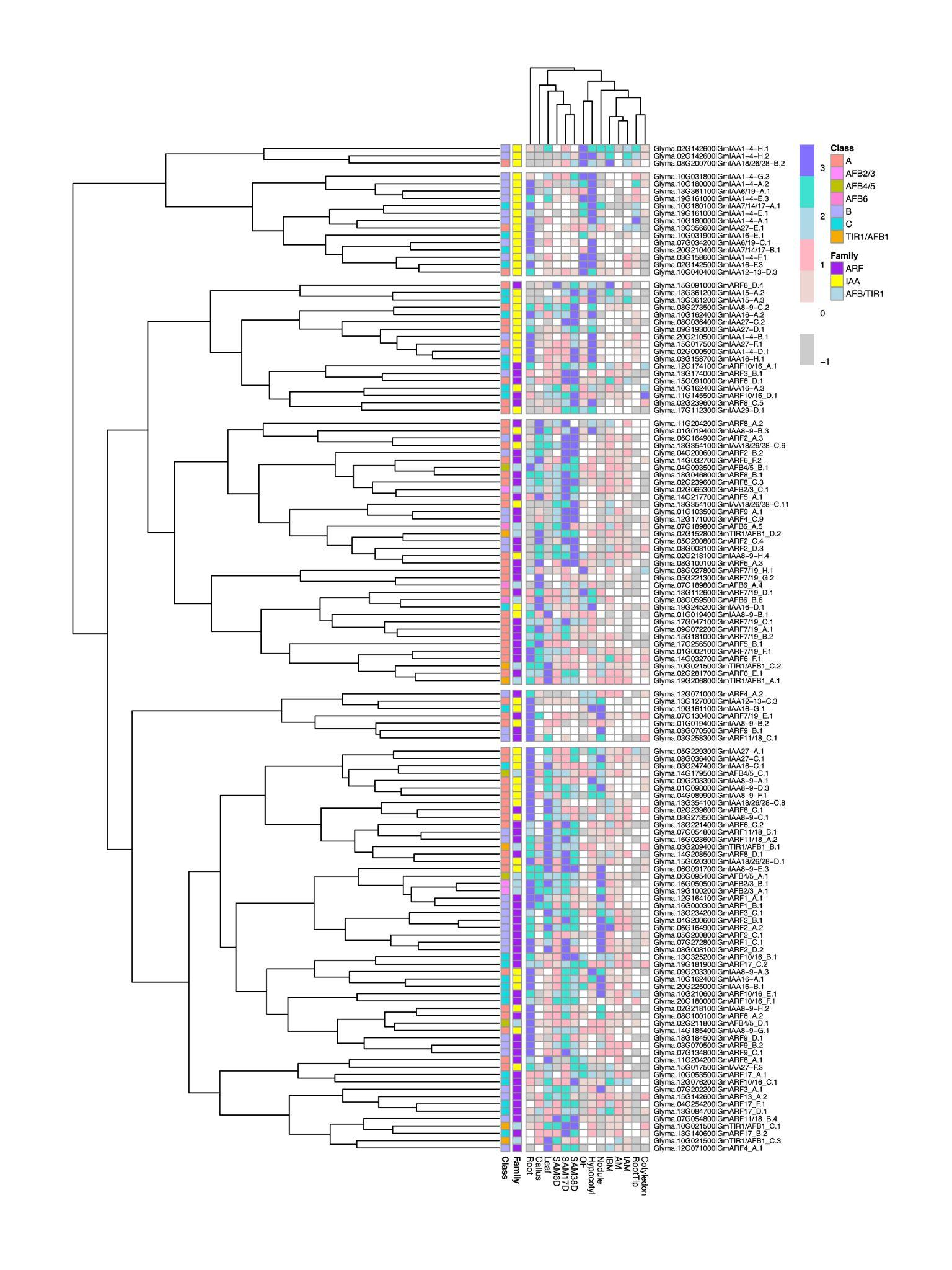

Figure S5. Heatmap of gene expression of G. max. Colors in the heatmap represent the z-score values of gene expression levels. The scale ranges from -1 in grey, representing gene expression levels below the mean of their respective rows. White color depicts values close to 0 or equal to 0, in which genes have expression levels similar to the mean expression of transcripts. Finally, positive values, ranging from 1 to 3, are depicted here in misty rose to blue and indicate gene expression levels above the mean expression of transcripts.

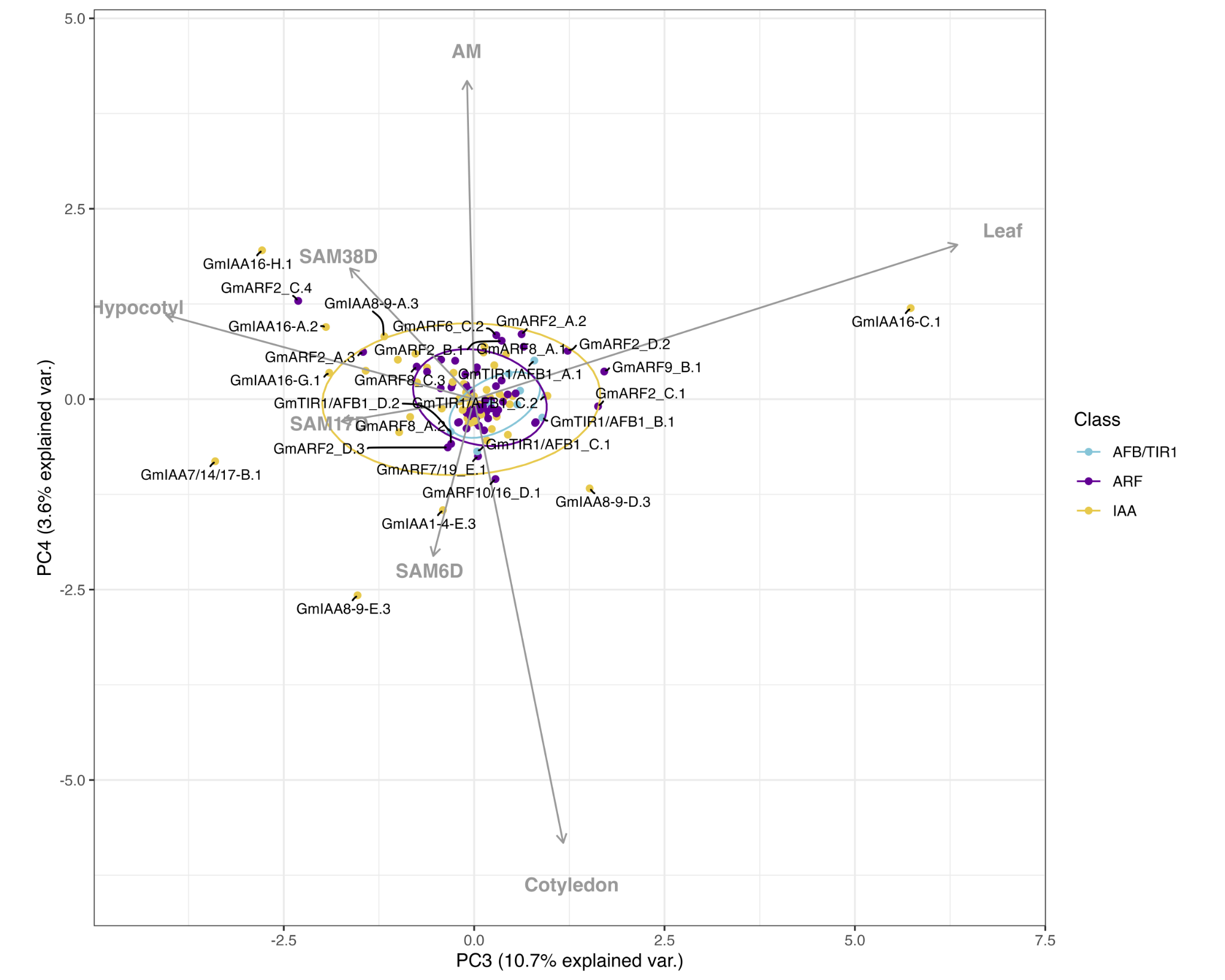

Figure S5. Correlation-based Principal Component Analysis (PCA): biplot of gene transcript expression and explanatory tissues involved in plant aerial architecture as eigenvectors (grey arrows), n = 7). Principal components 3 and 4 account for 14.3% of the total inertia.
